# How certain are we? Development of an ensemble based framework for assessing astronaut cancer risks from space radiation

**DOI:** 10.1101/2021.01.29.428854

**Authors:** Lisa C. Simonsen, Tony C. Slaba

## Abstract

A new approach to NASA space radiation risk modeling has successfully extended the current NASA probabilistic cancer risk model to an ensemble framework able to consider sub-model parameter uncertainty as well as model-form uncertainty associated with differing theoretical or empirical formalisms. Ensemble methodologies are already widely used in weather prediction, modeling of infectious disease outbreaks, and certain terrestrial radiation protection applications to better understand how uncertainty may influence risk decision-making. Applying ensemble methodologies to space radiation risk projections offers the potential to efficiently incorporate emerging research results, allow for the incorporation of future models, improve uncertainty quantification for underlying sub-models, and reduce the impact of subjective bias on risk projections. Moreover, risk forecasting across an ensemble of multiple predictive models can provide stakeholders additional information on risk acceptance if current health/medical standards cannot be met for future space exploration missions, such as human missions to Mars. In this work, ensemble risk projections implementing multiple sub-models of radiation quality, dose and dose-rate effectiveness factors, excess risk, and latency as ensemble members are presented. Initial consensus methods for ensemble model weights and correlations to account for individual model bias are discussed. In these analyses, the ensemble forecast compares well to results from NASA’s current operational cancer risk projection model used to assess permissible mission durations for astronauts. However, a large range of projected risk values are obtained at the upper 95^th^ confidence level where models must extrapolate beyond available biological data sets. Closer agreement is seen at the median + one sigma due to the inherent similarities in available models. Identification of potential new models, epidemiological data, and methods for statistical correlation between predictive ensemble members are discussed. Alternate ways of communicating risk and acceptable uncertainty with respect to NASA’s current permissible exposure limits are explored.

## II. Introduction

### II.I Challenges in space cancer risk projection

For future missions beyond low Earth orbit (LEO), astronauts exposed to space radiation are at increased risk of potential in-flight performance decrements and long-term health consequences including radiogenic cancers, cardiovascular disease (CVD), and possible insult to the central nervous system (CNS) resulting in cognitive or behavioral impairments. To protect astronauts from the risks associated with space radiation exposure, NASA has defined permissible exposure limits (PELs) [NASA 2014]. The current PEL for cancer is defined such that astronaut exposures do not exceed a 3% risk of exposure induced death (REID) evaluated at a 95% confidence level (CL) (equivalent to the 97.5^th^ percentile). This CL has been set to protect against significant uncertainties in the cancer risk projection associated with a lack of directly relevant experimental and epidemiological data for humans exposed to space radiation. Modeling plays a critical role in capturing our current state of knowledge explicitly expressed by evaluation of the CL. For design and mission planning, remaining below the cancer PEL is thought to minimize risk (or remain below possible thresholds) for other long-term health decrements (i.e., CVD, CNS).

Characterizing and communicating the space radiation risk landscape to a diverse group of stakeholders and decision-makers over a broad range of exploration mission architectures remains challenging due to the uncertainties involved and insufficient reliable data to fully anchor projections. Meeting today’s PELs can be difficult for crew with previous spaceflight experience and for young female crew selected for lunar surface missions. Mars mission architectures present even greater challenges with radiation risks that exceed NASA career PELs for all crew [Cucinotta et al. 2013]. In such cases where health or medical standards cannot be met or the level of knowledge does not permit a standard to be developed, NASA has established an ethical framework to accept increased risk [IOM 2014]. Health risks and associated uncertainties must be quantified to guide informed decision-making.

Inclusion of international crew and post-mission treatment requirements may impose further complexity. NASA’s Strategic Plan for Lunar Exploration [NASA 2020] will foster opportunities for international partnerships and most likely lead to international crews for lunar sustainability and Mars missions. The world’s space agencies that are currently involved in human spaceflight use a variety of methods to assess radiation exposure and risk to crew, as well as a variety of protection quantities and limits. Efforts are underway by the International Commission on Radiological Protection (ICRP) to develop a common health risk assessment framework and provide recommendations on exposure limits for exploration-class missions among the International Space Station partner countries [ICRP 2019]. With the implementation of the TREAT (To Research, Evaluate, Assess, and Treat) Astronauts Act (https://www.congress.gov/bill/114th-congress/house-bill/6076), NASA will provide former astronauts and payload specialists with monitoring and treatment for psychological and medical conditions associated with spaceflight. This new Act may necessitate probabilistic assessments at higher confidence intervals and/or additional applied and translational research data sets (e.g. mutational signatures) to understand whether conditions can be identified as associated with spaceflight. Assessments at these extreme confidence intervals are particularly sensitive to the underlying model assumptions used to represent biological outcomes, and there exists a general lack of available data with sufficient statistical quality to substantively resolve these uncertainties in risk model projections.

The NASA Space Cancer Risk model translates human epidemiology data from an acutely exposed 1940s Japanese population to a present day US healthy population (e.g. astronauts) chronically exposed to space radiation. Here, we refer to NSCR2020 as a “single model” comprised of distinct sub-models selected for each of the major components of Fig 1. For certification with the NASA PEL, risk must be evaluated probabilistically, thus requiring uncertainties to be defined for each of these components. Uncertainties have been assigned to underlying sub-model parameters based on available epidemiological data, limited experimental data and/or subject matter expert opinion. Other inherent uncertainties within this model framework exist in the applicability of scaling risk from a terrestrial population acutely exposed to predominantly gamma radiation (left hand side of Fig 1) to an interplanetary crew exposed to a vastly different extraterrestrial space radiation environment (right hand side of Fig 1) that are not quantified in current sub-model parameterizations (See Section III.II). The CL applied in the cancer PEL is thought to protect against these uncertainties as well.

**Fig 1.**
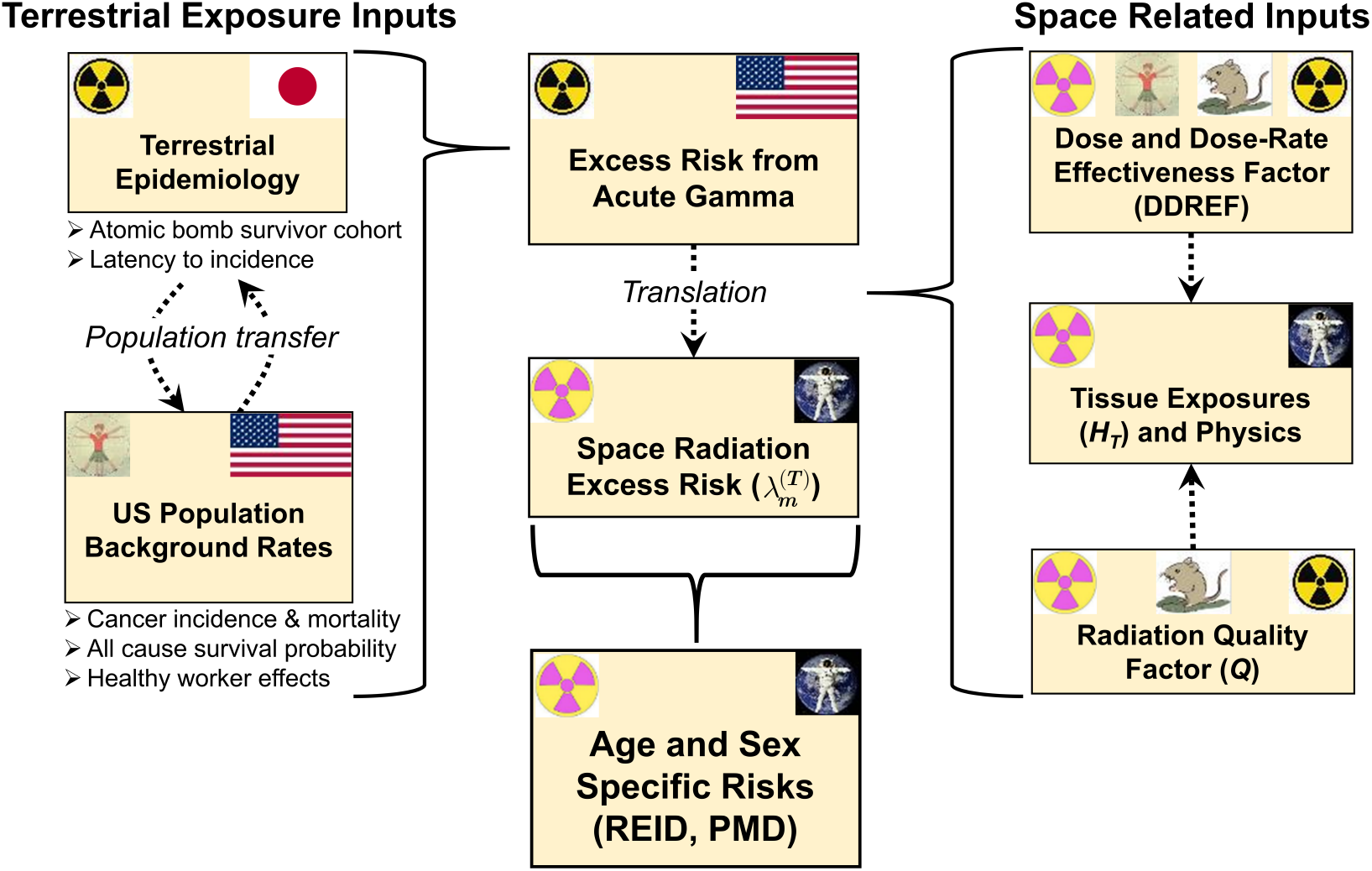
Notional implementation of the NASA’s current space radiation cancer risk model illustrating the use of epidemiological and radiobiology data sets to scale cancer incidence and mortality in exposed terrestrial populations to space-based estimates of radiogenic cancer risk.

Although significant new space-relevant data sets are not available, re-analysis of existing data is ongoing, and additional models continue to be developed to describe radiation quality [Cucinotta 2015], dose-rate effects [Kocher et al. 2018; Cucinotta and Cacao 2017], and dose response in the atomic bomb survivor cohort [Kaiser et al. 2012; Kaiser and Walsh 2013]. A strategy on how or when to incorporate newly developed state-of-the-art models into NASA risk assessments is critical, since experience has shown that such advancements can often lead to noticeably different REID projections at the upper 95% CL [Cucinotta and Cacao 2017] but often without sufficient data to anchor or independently validate such changes [Chappell et al. 2020]. Such dynamic changes to underlying sub-models and risk projections can be problematic for operational planning as well as mission architecture design.

Radiation exposure estimates and risk projections depend on multiple factors such as mission destination and duration, vehicle design, and heliospheric conditions. For illustrative purposes, analyses presented here are based on a ~6 month extended lunar orbital design reference mission (DRM) (https://www.nasa.gov/topics/moon-to-mars/lunar-gateway) during solar minimum conditions (time of maximum GCR flux) for a 35-year-old female astronaut (never smoker) behind nominal spacecraft shielding of 20 g/cm^2^ aluminum^1^. Specifics of the DRM were selected such that the upper 95% CL REID value, denoted as *R*_95%_, is aligned with the 3% PEL (Fig 2) for direct comparison with NASA’s current limit. The average effective dose [ICRP 2007] to females was found to be approximately 185 mSv. Utilization of male tissue shielding models resulted in exposures differences of <3%. External body exposures from spacecraft shielding alone (i.e. just outside the astronaut) are slightly higher, approximately 82 mGy and 215 mSv, and are consistent with previous measurements [Zeitlin et al. 2013] and model assessments [Simonsen et al. 2020]. The resulting probabilistic REID distribution from NSCR2020 is shown in Fig 2 along with the median (*R*_med_) and upper 95% CL (*R*_95%_) values.

**Fig 2.**
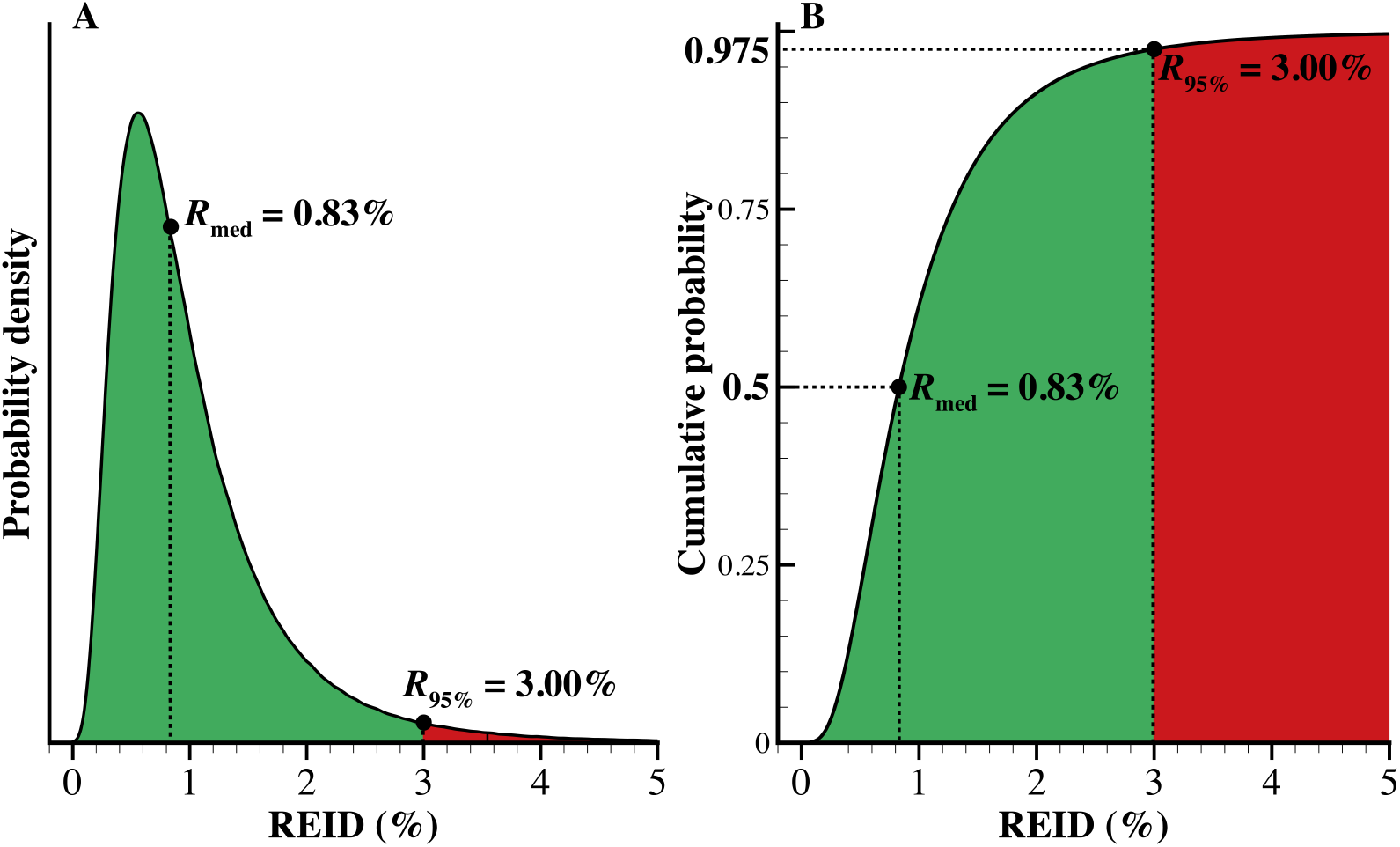
Probability density (A) and cumulative probability (B) functions of REID (radiation exposure induced death) for a 35-year-old female astronaut (never smoker) on a 6 month mission behind 20 g/cm^2^ of aluminum shielding during solar minimum conditions. The Badhwar-O’Neill 2020 GCR model [Slaba and Whitman 2020] combined with the HZETRN radiation transport code [Slaba et al. 2020] are used to evaluate relevant physical quantities. The median (*R*_med_) and upper 95% CL (*R*_95%_) values are explicitly shown. Specifics of the DRM were selected such that *R*_95%_ compares directly with the 3% PEL. The probability density function (A) is normalized so that the area under the curve is unity; the area of the red shaded region is 0.025.

A fold-factor is often computed as (*R*_95%_/*R*_med_) to assess overall uncertainty of the risk projection. For the DRM configuration of Fig 2, the NSCR2020 fold factor is 3.6 and reflects the fact that biological effects of the primary and secondary space radiation environment seen by astronauts are poorly understood. Differences in the temporal and spatial deposition of energy in tissues from space radiation impart unique biological damage to biomolecules and cells compared with terrestrial radiation (such as gamma radiation), which, for a given dose is much more damaging. As shown in Fig 1, models of radiation quality and dose and dose-rate effectiveness (DDREF) are employed to translate epidemiological data of terrestrial exposures to space-based risks. The current risk acceptance paradigm, as defined by the PEL and single model risk projection, is highly sensitive to the low-probability tails of one or more of the underlying sub-model uncertainties (e.g., radiation quality). The data to support these low probability tails are extremely limited even compared to the already sparse datasets to which more probable risk projections are anchored. In such situations, extreme model uncertainties become increasingly dependent on initial assumptions (or model form) and subjective decisions that cannot be robustly tested or validated with available data.

To illustrate, consider the radiation quality component of the risk model (Fig 1) which characterizes increased relative biological effectiveness of the particles and energies comprising the space environment compared with gamma radiation. The NASA quality factor, *Q*(*E*,*Z*) [Cucinotta et al. 2013], is described by a biophysical model calibrated to available animal and cellular experimental data. Included in this model is the parameter, *κ*, related to the ion and energy of maximal biological effectiveness. Cucinotta and colleagues [2013] identified preferred *κ* values for light ions and heavy ions based on the limited experimental data available. Uncertainty distributions were then subjectively assigned to *κ* for use in probabilistic risk projections (as implemented in NASA’s previous 2012 risk model). Subsequent updates to the quality factor model [Cucinotta 2015, Cucinotta and Cacao 2017] followed a similar approach. In these publications, the relative uncertainties (with respect to the point estimates) for *κ* were subjectively assigned as normally distributed with a mean of one and a standard deviation of one-third. NASA’s current operational model, NSCR2020, now implements the relative uncertainties for *κ* as log-normally distributed (with same mean and standard deviation) to ensure maximal biological effectiveness is assigned to ions and energies in a more realistic manner [Simonsen and Slaba 2020]. This seemingly minor change to a single parameter uncertainty distribution is found to have a significant impact on risk projections at large confidence levels (Fig 3). Of particular interest are the values of *κ*, shown in red, contributing to the upper tail of the REID distribution. The *R*_med_ values from both models are identical since they are both reasonably anchored by available experimental data. However, the *R*_95%_ value from NSCR2012 is 20% higher and is driven by low probability *κ* values characterized by the subjective selection of a normal versus log-normal uncertainty distribution. Sparsity of relevant data precludes a more robust and objective characterization for parameter uncertainties in this case. Here, the *R*_95%_ value is shown to be acutely sensitive to subjective assumptions and decisions included in sub-model parameters.

**Fig 3.**
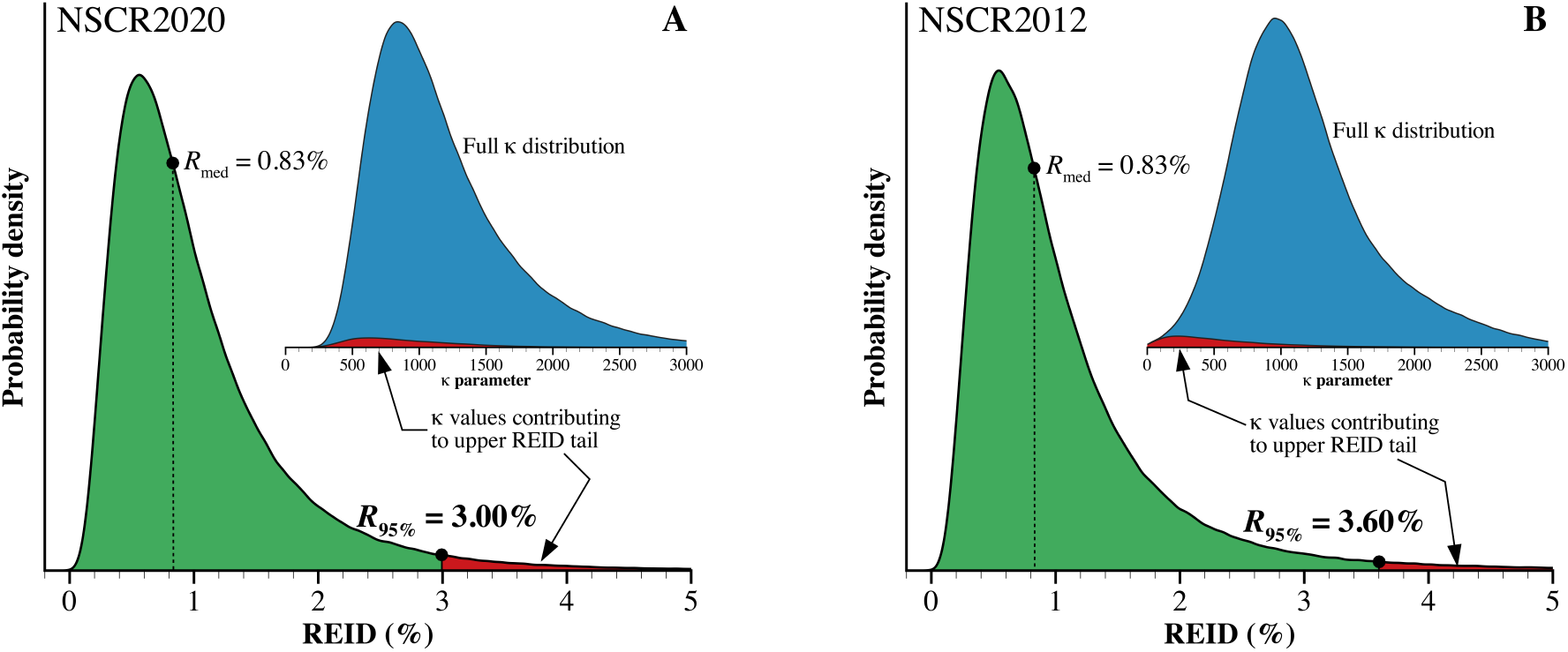
Calculated REID distributions from both NSCR2020 and NSCR2012 with related information about *κ* in the insets illustrating the sensitivity of the upper 95% CL of REID (*R*_95%_) to a specific parameter uncertainty distribution. (A) Current NASA model (NSCR2020) where the quality factor parameter, *κ*, is described by a lognormal distribution. (B) Previous NASA model (NSCR2012) where *κ* is described by a normal distribution. Results are for a 35-year-old female astronaut (never smoker) on a 6 month mission behind 20 g/cm^2^ of aluminum shielding during solar minimum conditions.

Likewise, median risk projection values, *R*_med_, have also been found to be sensitive to these kind of modeling assumptions. Recent publications by Cucinotta and colleagues [Cucinotta and Cacao 2017, Cucinotta et al. 2020a] examine the impact of non-targeted effects (NTE) in representing low dose biological responses that appear as modifications to the quality factor component of the risk model. In their analysis, *R*_med_ values were calculated separately using both a targeted effects model that is similar to what is used in NSCR2020 as well as a NTE model with different bystander effect sizes (number of cells nearby a directly hit cell that receive an intercellular, potentially carcinogenic signal). The *R*_med_ estimates produced by the targeted and non-targeted effects models differed by factors of two or more depending on the bystander effect size chosen. While the existence and relevance of NTE effects to risk have been shown [Barcellos-Hoff and Mao 2016], available data sets to sufficiently resolve model assumptions remain elusive [Chappell et al. 2020, Cucinotta and Cacao 2017].

### II.II A new approach for space cancer risk projection

Ensemble modeling offers a different approach in assessing uncertainties and represents a paradigm shift from NASA’s current methodology of implementing a single model. Here, the goal is *not* to identify the *“best”* model but rather to use information from multiple single models or underlying sub-models to better describe risk when faced with large uncertainties and limited data. Over the past decade, the National Hurricane Center has greatly improved its forecasting by relying on consensus forecast models using various simple and weighted combinations of predictive models [Slingo and Palmer 2011]. Other entities, including the World Health Organization have considered ensemble approaches to predict the health impact of vaccines on the transmission of infectious diseases, such as malaria, using multiple predictive models where immunity and variability in host response are poorly understood [Smith et al. 2012]. Likewise, the annual prediction of influenza season severity and timing has been modeled using weighted-density ensemble methodologies to obtain single predictions that leverage the strengths of each ensemble model member [Ray and Reich 2018]. New to space weather applications, researchers are also looking toward ensemble forecasting as a way to combine the multiple predictive models being developed for predicting solar flare activity by linearly combining the probabilistic forecasts from a group of operational forecasting methods [Guerra et al. 2020].

More directly relevant to space radiation cancer risk projection, the Oak Ridge Center for Risk Analysis (ORCRA) recently developed an ensemble DDREF model [Kocher et al. 2018] which will be explicitly considered here. Other statistical approaches to assess risk from radiation exposure include multi-model inference (MMI) analyses to develop a joint risk estimate from several plausible models rather than relying on a single model of choice. MMI statistical analyses employing multiple biologically-based and/or empirical models have been used to model the radiogenic risks of leukemia, breast cancer, cerebrovascular, and heart disease using Japanese atomic bomb survivor data [Kaiser et al. 2012, Kaiser and Walsh 2013, Schöllnberger et. al 2018]. The analyses of Kaiser and colleagues [Kaiser et al. 2012, Kaiser and Walsh 2013] have shown the potential to produce more reliable point estimates, reduce bias, and provide a more comprehensive characterization of uncertainties of radiogenic risks.

Here, we describe the development of alternate methods of evaluating risk for US exploration missions through a statistical analysis using the ensemble of sub-models described in Fig 1. This approach can facilitate the comparison of NSCR2020 with historical models of risk [ICRP 1991], alternate models [Cucinotta et al. 2020a], and future international models [Walsh et al. 2019] which may prove important in planning for international exploration crews. Moving to an ensemble framework provides an opportunity to shift the focal point of astronaut risk projection from a region of uncertainty, sensitivity, and subjective bias (the *R*_95%_) toward regions of stability and general agreement between models more directly determined by the bulk of experimental and epidemiological data. While ensemble methods can aid in assessing uncertainties in extrapolation beyond these limited data sets, significant new data sets are still needed to improve model development and appreciably reduce uncertainties in astronaut risk projection. Risk forecasting across an ensemble of multiple predictive models can provide crew and decision makers a comprehensive assessment of current and future mission risk landscapes, especially when mission architectures may not immediately meet established PELs and decisions are needed for the informed acceptance of additional risk.

## III. Materials and Methods

### III.I NASA exposure limits and current risk modeling

#### III.I.I Space radiation permissible exposure limits

Permissible limits are currently defined for low Earth orbit (e.g. International Space Station missions), such that “planned career exposure to ionizing radiation shall not exceed 3 percent REID for cancer mortality at a 95 percent confidence level to limit the cumulative effective dose (in units of Sievert) received by an astronaut throughout his or her career [NASA 2014].” Unlike terrestrial dose limits, where risk as a function of dose can often be more reasonably estimated from response to low-LET^2^ irradiations, NASA has established their limits in terms of risk to account for the complex nature of HZE radiation in space where equal absorbed doses of space radiation and terrestrial radiation do not have the same biological effects. A 95% confidence level has been included in the limit to account for large uncertainties in risk projections including the understanding of the radiobiology of heavy ions, dose-rate and dose protraction and limitations in human epidemiology data. This confidence interval is the basis for establishing allowable cumulative effective dose (Sievert, Sv) and consequently a crew member’s permissible mission duration (PMD) based on previous and projected exposures. By establishing career limits in terms of risk, rather than dose, it was anticipated that NASA could modify corresponding dose limits (Sv) to allow for increased PMDs as scientific knowledge describing the relationship between risk and dose evolved and uncertainties were reduced [NCRP 2014]. Following established processes within NASA’s Office of the Chief Health and Medical Officer (OCHMO), scheduled reviews of the health standards are conducted every five years [NASA 2016].

#### III.I.II Current NASA risk model

The National Council on Radiation Protection (NCRP) report No. 132, “Radiation Protection Guidance for Activities in Low-Earth Orbit [NCRP 2000],” forms the basis of NASA’s radiation protection approach [NASA 2014] and the majority of methodologies implemented in NASA’s cancer risk projection model. NSCR2012 [Cucinotta et al. 2013] was a significant step forward in gathering and making use of the available radiobiology and epidemiology data into a single formalism capable of projecting astronaut cancer risk for past and future missions. The current operational version, NSCR2020, used by NASA to certify crew for missions has been significantly updated over the years to include recommendations from a National Academy of Science review in 2012 [NRC 2012] as well as other corrections to underlying models and software [Slaba et al. 2010, 2020; Slaba and Whitman 2020]. The model estimates probabilistic REID in accordance with the NASA PEL. This REID quantifies the lifetime mortality risk attributable to radiation exposure and accounts for competing causes of death. It is calculated by folding a tissue-specific mortality rate, 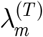, against a hazard function and integrating over age according to

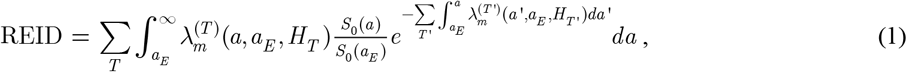

where *a_E_* is the age at exposure, *a* is the attained age, *S*_0_ is the survival probability for the unexposed background population [Arias 2015], and the summation is taken over all radiosensitive tissue types, *T*. The variables *a*’ and *T*’ are integration variables with the same meaning as *a* and *T*, respectively.

For solid cancers, the sex- and tissue-specific radiation-induced mortality rate, 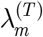, is calculated in terms of the corresponding incidence rate, 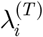, as

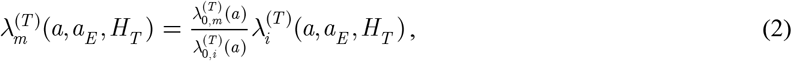

where 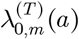 and 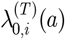 are the sex- and tissue-specific cancer mortality and incidence rates for the background population of interest, respectively. The radiation-induced solid cancer incidence rate is written here as

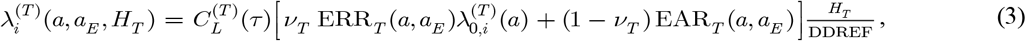

where ERR_*T*_ and EAR_*T*_ are the sex- and tissue-specific excess relative and absolute excess risk functions, respectively, and *v_T_* is the transfer weight defining the contributions of the relative and absolute risks to the total. The term 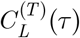, with *τ* = *a* – *a_E_*, is the latency factor used to describe the time-lag between age at exposure and first appearance of cancer.

Combining equations (2) and (3) yields the final form for the mortality rate

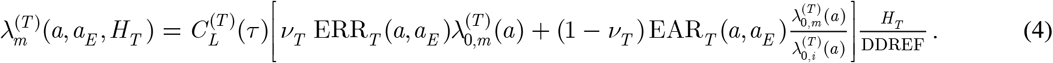

For leukemia, the ERR and EAR are derived directly from mortality data, and the scaling of equation (2) is not needed. The tissue dose equivalent, *H_T_*, is computed as

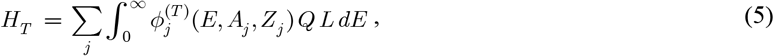

Where 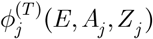 is the fluence of type *j* particles with atomic mass *A_j_* and charge number *Z_j_*, *Q* is the quality factor, and *L* is the LET.

The DDREF and *Q* are the scaling factors specifically used to translate the radiation-induced cancer mortality rate derived from populations exposed to acute gamma radiation to the low dose-rate and mixed-LET exposure characteristic of the space environment. The quality factor has been defined thus far by fitting parametric biophysical models to available relative biological effectiveness (RBE) data ascertained from experiments with animals and cells for various biological endpoints. The DDREF is derived from a combination of radiobiological studies and epidemiological data. The instantaneous, low-LET excess risk terms are derived from the Life Span Study (LSS) of the atomic bomb survivor cohort assuming a linear no-threshold (LNT) dose response; it includes both absolute and relative risk parameterizations [NRC 2006, UNSCEAR 2006, Preston et al. 2007, Little et al. 2008] to transfer risks to a United States background population with or without a history of smoking. As noted by Cucinotta and colleagues [2013] and in the National Research Council (NRC) review of NSCR2012 [NRC 2012], uncertainties associated with *Q*, DDREF, and excess risk terms remain the major sources of uncertainty currently accounted for in the model. As will be discussed, other sources of uncertainty not accounted for in the model formalism may also be significant [Cucinotta et al. 2013, NCRP 2014].

In order to ensure that crew career limits are not exceeded, for planned mission exposures, equation (1) must be evaluated probabilistically to account for uncertainties associated with the various parameters and assumptions contained within the radiation mortality and survival terms. The quantities needed to evaluate equation (1) such as *Q*, DDREF, and the excess risk functions have varying levels of uncertainty arising from either a lack of relevant data and/or knowledge, inherent variability that cannot be reduced with new data, or both. These uncertainties are propagated into REID assessments so that critical values such as *R*_med_ or *R*_95%_, can be obtained. As in NSCR2012, Monte Carlo procedures are used to perform this probabilistic analysis, requiring uncertainty distributions for each of the pertinent values and assumptions contained within equation (1). These distributions, intended to represent the range of possible parameter values, are either assigned through subjective assessments of available data or objectively determined through uncertainty quantification when enough data exist. The uncertainty distributions included in the NASA model are summarized in Table 1.

**Table 1.**
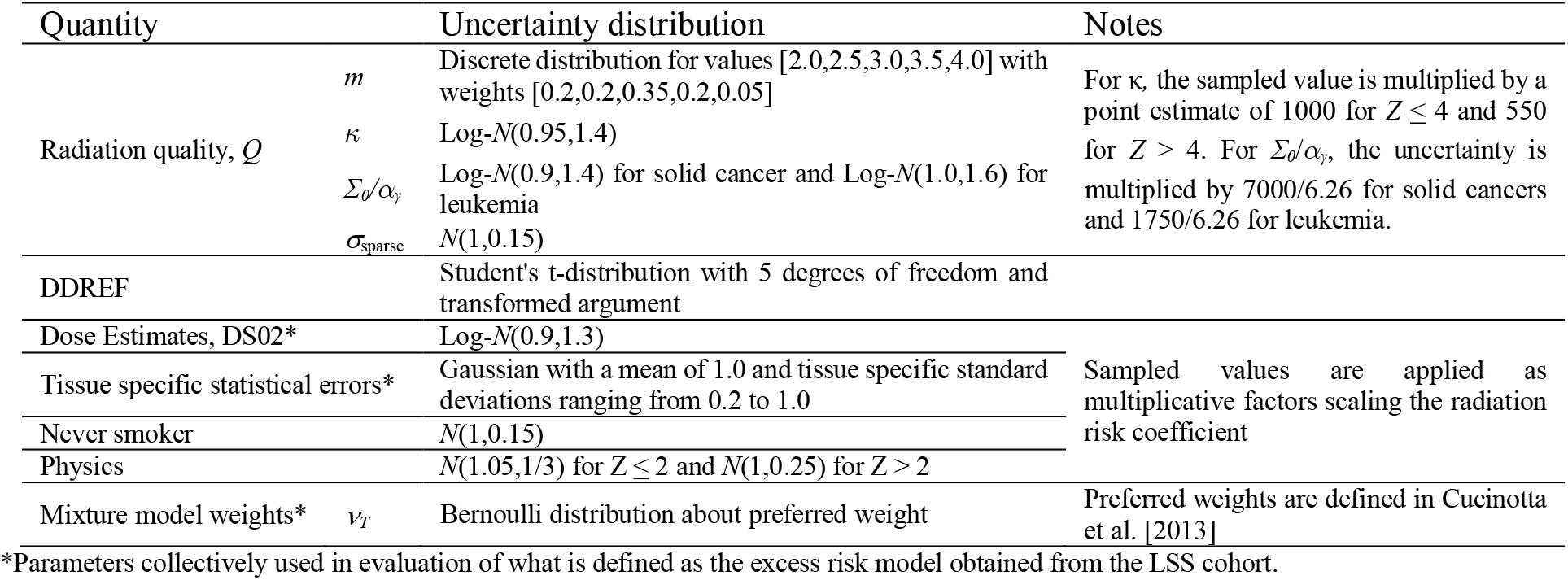
Summary of uncertainties used in NSCR2020. *N*(*μ*,*σ*) refers to the normal distribution with mean *μ* and standard deviation *σ*. Log-*N*(*μ_G_*,*σ_G_*) refers to the log-normal distribution with geometric mean *μ_G_* and geometric standard deviation *σ_G_*.

To understand the relative impact of the major uncertainties on the final REID assessment, sensitivity tests can be performed wherein parameters of interest are sampled probabilistically while all other parameters are held fixed at their preferred values, or point estimates. Results from this type of sensitivity analysis are given in Fig 4 for the extended lunar orbital DRM mission configuration. The items identified in Table 1 as DS02, tissue specific statistical errors, and mixture model weights are collectively identified in Fig 4 as the excess risk model. These values and parameters are derived from the LSS cohort [Kotaro et al. 2012] to describe cancer incidence and mortality risk following acute exposure to low-LET radiation. Included in this model is the transfer of risk to the appropriate population of interest as defined by equations (2) – (4). The DDREF and *Q* are then used to scale these low-LET risks to the mixed LET environment in space. It can be seen in Fig 4 that uncertainty associated with *Q* is the dominant term contributing to the total fold-factor of 3.6. Other sources of uncertainty not accounted for in the model formalism may also be significant [Cucinotta et al. 2013, NCRP 2014] (see Section III.II).

**Fig 4.**
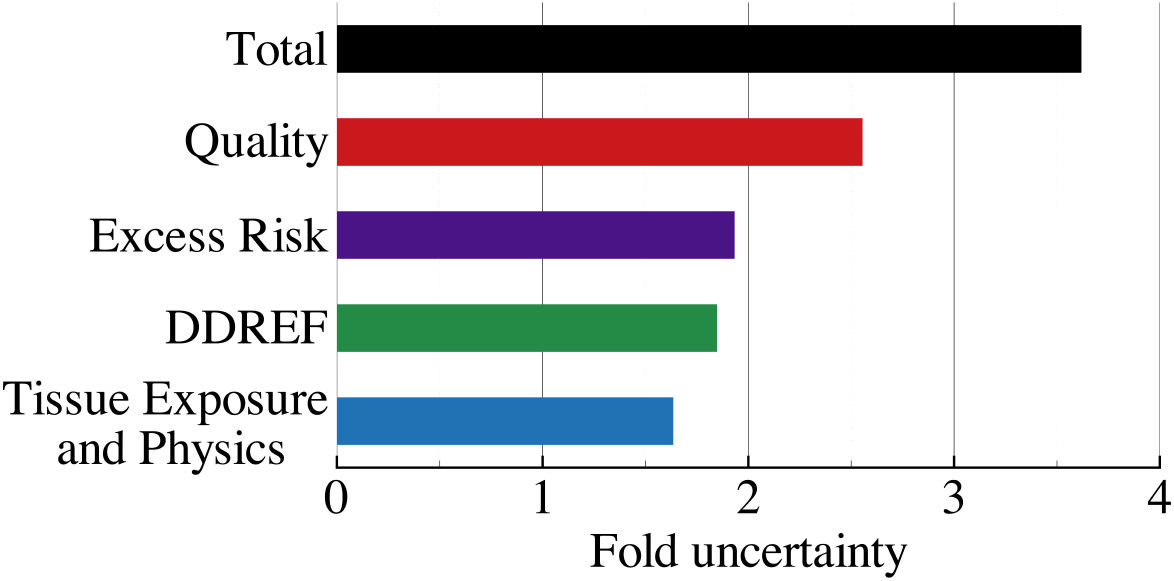
NSCR2020 uncertainty in probabilistic REID assessment associated with radiation quality, excess risk, dose and dose-rate effectiveness factor (DDREF), and physics for an extended lunar orbital design reference mission. REID, radiation exposure induced death.

#### III.I.III Permissible mission durations

Permissible mission durations (PMD) for individual crew members may be calculated from equation (1) by determining the mission exposure (and hence, duration) required to yield a *R*_95%_ = 3% for a given age at exposure. For the exposure ranges of interest to human missions, REID is nearly directly proportional to effective dose (Sv) which accumulates with mission duration. For a given mission configuration with duration *d_m_* and projected *R*_95%_ value, the PMD for an astronaut to remain in the radiation environment characterized for the mission may be calculated as PMD = 3*d_m_*/*R*_95%_. PMD are directly dependent on the projected space radiation environment and thus, time in solar cycle. Within our solar system, the solar wind modulates the flux of GCR over an approximate 11-year cycle with an intensity that is inversely correlated with solar activity. During phases of higher solar activity, the GCR intensity is at a minimum, whereas at solar minimum, the GCR intensity is maximal. At solar maximum, effective dose estimates behind typical spacecraft shielding are reduced by roughly a factor of 2 compared with solar minimum dose estimates [Townsend et.al 1990, Slaba et. al 2016]. Calculations in the deep-space environment are used to guide long-term mission design and are shown in Table 2 for both solar minimum and maximum solar conditions. These values can be compared with mission durations required for one year deep space habitat (365 days) and Mars short stay (621 days) exploration-class missions.

**Table 2.** Permissible mission durations beyond low Earth orbit as a function of age and sex for crew with no previous radiation exposure. Calculations were performed using NSCR2020 for never smokers with 20 g/cm^2^ of aluminum shielding during both solar minimum (June 1976) and solar maximum (June 2001) conditions. Average exposure rates external to the body are provided along with the effective dose rate (mSv/day). Only minor differences (<3%) were found as a result of tissue shielding between males and females.

| Age at exposure (years) | Permissible mission duration (days) |  |  |  |
| --- | --- | --- | --- | --- |
|  | Solar minimum |  | Solar maximum |  |
|  | Female | Male | Female | Male |
| 30 | 163 | 236 | 312 | 453 |
| 35 | 174 | 248 | 336 | 477 |
| 40 | 186 | 262 | 357 | 503 |
| 45 | 198 | 276 | 381 | 530 |
| 50 | 210 | 291 | 407 | 560 |
| 55 | 225 | 311 | 435 | 596 |
| 60 | 243 | 335 | 470 | 645 |
| Dose (mGy/day) | 0.47 |  | 0.23 |  |
| Dose eq. (mSv/day) | 1.24 |  | 0.65 |  |
| Effective dose (mSv/day) | 1.06 |  | 0.56 |  |

### III.II Ensemble-based methods for radiogenic cancer risk prediction

Consensus projections formulated on the basis of perturbed parameter (PP) and multi-model (MM) methods, on average, are more accurate than predictions from their individual model or sub-model components [Fritsch et al. 2000; Ray and Reich 2018]. Although it has not previously been described in these terms, the NSCR2020 model employs PP methods which are an aspect of ensemble forecasting. In PP schemes, uncertain parameters within a specific model are repeatedly perturbed to yield distinct outcomes which collectively form a distribution. Parameter perturbations may be guided by random noise, subject matter expert opinion, or direct comparison of the parameter to measured data, if possible. As noted, the evaluation of REID employs a series of sub-models which are parametric in nature with uncertainty distributions assigned to each of the relevant parameters (Table 2). Monte Carlo methods are then used to randomly sample parameters within the prescribed distributions over numerous trials to calculate a probabilistic distribution of REID values. Given a large enough sampling of perturbed parameters, the PP approach appears similar to the Monte Carlo uncertainty propagation methods employed in NSCR2020.

While PP schemes account for parameter uncertainty, they do not capture uncertainty associated with the fundamental assumptions or form of an underlying model [Tebaldi and Knutti 2007]. This additional source of uncertainty is sometimes referred to as model-form error and may be addressed with multi-model (MM) ensemble forecasting. In MM methods, multiple predictive models that may be based on different theoretical formulations, assumptions, data sources, or solution methodologies are evaluated as ensemble members. In applications where basic principles and mechanisms are well understood and significant data exist, PP and MM schemes may not provide significantly different information about overall model uncertainties. However, astronaut risk projection relies on sparse experimental datasets in cells and animals for which the driving mechanisms of carcinogenesis are not fully understood and methods for translating available experimental data to astronauts in the space environment remain highly uncertain. Thus, even as individual sub-models (e.g., radiation quality or dose-rate effects) are developed and improved, PP ensemble results would only partially reflect uncertainties in risk projection. MM methods may offer an additional avenue to better characterize the current state of scientific knowledge and the associated uncertainties.

Fig 5 illustrates the computational framework considered here for MM ensemble forecasting of astronaut risk. The green boxes denote the sub-models specifically used in this analysis and the red boxes indicate emerging models or epidemiology studies that can be incorporated in future assessments. Although, there are a limited number of sub-models available which incorporate non-trivial differences in underlying assumptions, functional forms, and/or quantified uncertainties, several distinct models do exist to demonstrate MM methods. For example, NSCR2020 and UNLV2017 implement distinctly different assumptions in quality and dose rate effects. The radiation quality factor used in NSCR2020 is based on the Katz biological risk cross section coupled with an initiation-promotion model of tumor induction [Katz et al. 1971, Wilson et al. 1993] and is calibrated against experimental RBE_max_ values. Recent updates to this radiation quality model (denoted as UNLV2017 box in Fig 5) consider separately a densely ionizing component associated with the ion track core and a sparsely ionizing component associated with longer range *δ*-ray electrons in the penumbra region [Cucinotta 2015]. Included in this separation is an important additional assumption that dose-rate effects are negligible in the track core and only appear in the sparsely ionizing *δ*-ray component. This modified radiation quality model is also calibrated against distinct experimental data, RBE_γ-acute_, instead of RBE_max_, resulting in a different uncertainty assessment than its NSCR2012 predecessor and with significant consequences on the upper 95% CL REID value. Likewise, several DDREF models [NRC 2006, Cucinotta et al. 2013, Cucinotta et al. 2017, Kocher et al. 2018] are readily available in the literature with non-trivial differences in model formulation and uncertainty quantification (green boxes in Fig 5). In establishing the initial ensemble framework, we have limited our consideration of ER models to those relying on the assumption of LNT dose response and epidemiological data from the LSS cohort, namely those included in NSCR2020 [BEIRVII-NRC 2006, UNSCEAR 2006, Preston et al. 2007, Little et al. 2008] and RadRAT [de Gonzalez 2012]. Aside from the uncertainties, which are included in RadRAT and excluded in NSCR2020, the latency models implemented here appear qualitatively similar with some non-trivial differences for leukemia and solid cancer [NRC 2006, de Gonzalez 2012]. The specifics of these sub-models are described in Simonsen and Slaba [2020]. While the current number of sub-models and distinct single models may be somewhat limited, numerous development activities are underway to provide additional data sets and methods for inclusion given similar endpoints of REIC (risk of exposure induced cancer) or REID (red boxes of Fig 5) (Section V.III Future ensemble members).

**Fig 5.**
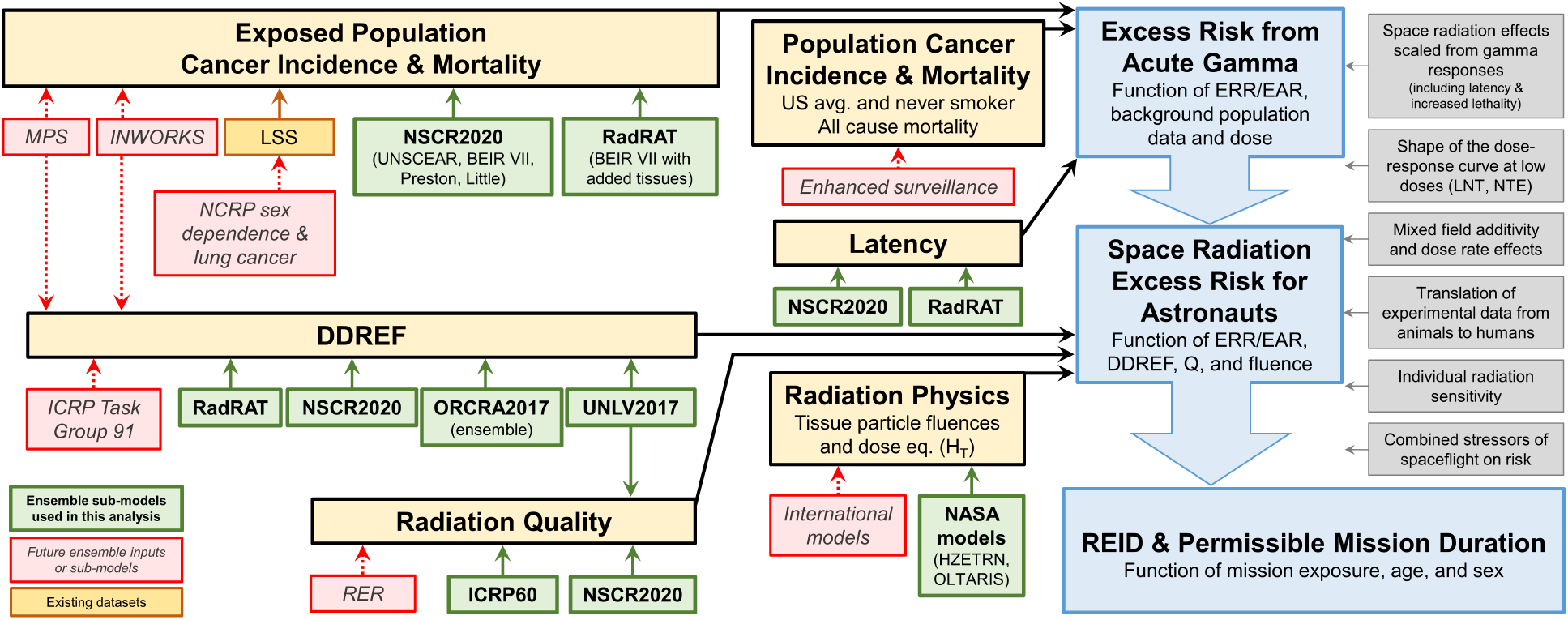
Example of current and near-term modeling and epidemiology studies that support ensemble methods in estimating the risk of cancer in astronauts from exposure to the space environment. Green boxes identify sub-models specifically considered in this analysis. Red boxes identify new models and data available in the near term. Gray boxes identify known uncertainties not explicitly accounted for in current risk model.

Additional known uncertainties, that are not explicitly accounted for, exist in the current risk projections (gray boxes of Fig 5) including assumptions about: the applicability of scaling space radiation effects directly from gamma responses resulting in a similar spectrum of tumors; the shape of the dose response curve at space relevant doses; the simple additivity of mixed field responses; the translation of exposed animal cohort data to humans; and the effects of individual sensitivity on projected risk. In addition to the space radiation environment, crew are exposed to a multitude of spaceflight stressors (such as, micro-gravity, isolation, confinement, and sleep disturbances) which may synergistically effect the actual risk. These uncertainties not only exist for projections of radiogenic cancers, but also for other long-term health effects associated with exposure to space radiation including central nervous system effects resulting in potential in-mission cognitive or behavioral impairment and/or late neurological disorders, degenerative tissue effects including circulatory and heart disease, as well as potential immune system decrements impacting multiple aspects of crew health. For long durations in space, the interdependency of these disease risk factors and combined stressors to the human as a system is largely unknown. Additional data in these areas will be required to inform the development of new models for future incorporation into an ensemble framework.

## IV. Results

### IV.I Comparison of ensemble risk projection with NSCR2020

Combining the multiple sub-models of radiation quality, DDREF, latency, and excess risk identified in Fig 5 provides a hybrid PP/MM ensemble risk projection tool where each sub-model contains its own description of parameter uncertainty and the combination of multiple models provides a framework to begin quantifying model-form uncertainty. While a larger number of relevant sub-models exist, here we focus on a limited number to extend current risk prediction methods to an ensemble framework. Fig 6 notionally illustrates how the various sub-models are combined.

**Fig 6.**
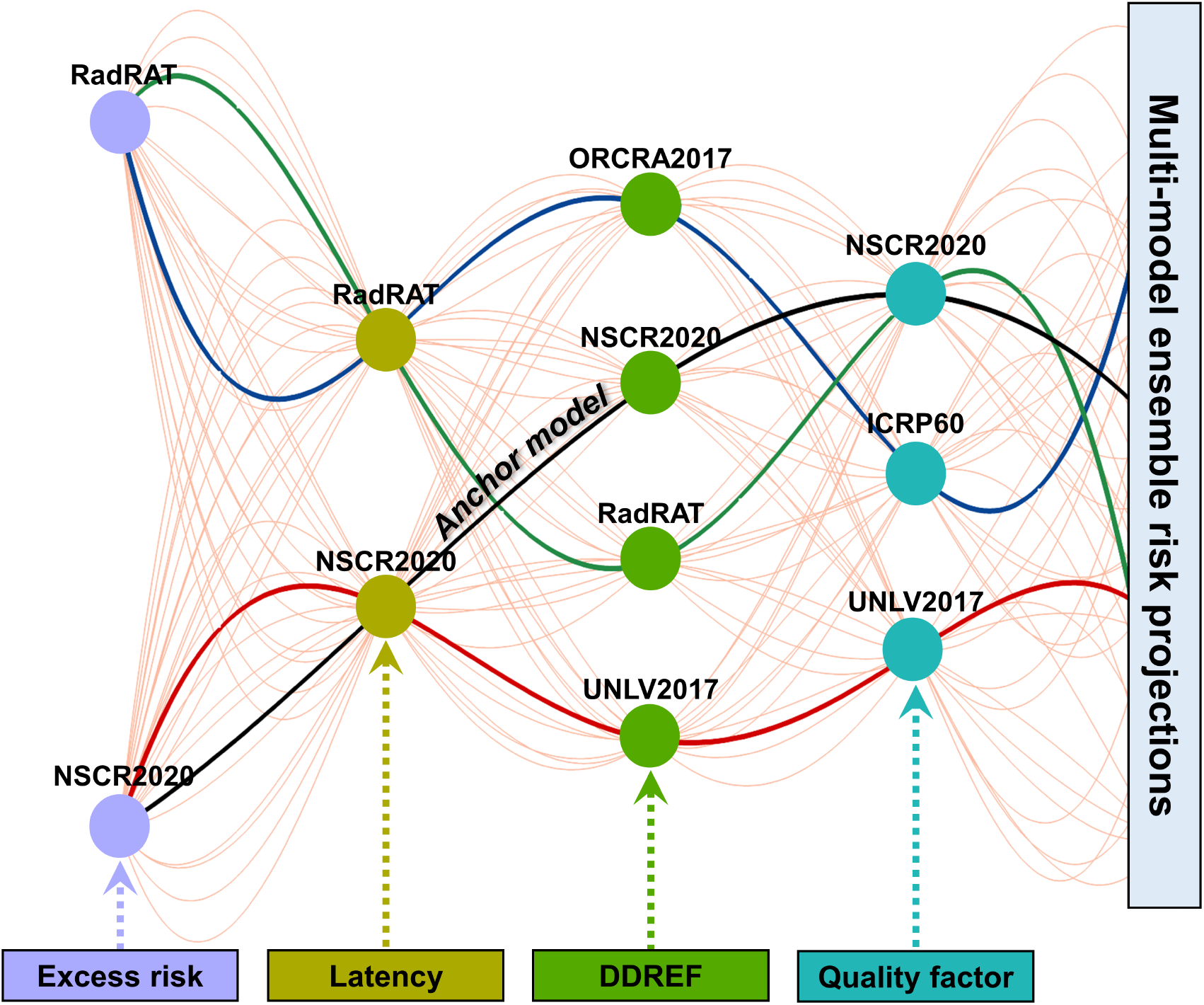
Visual depiction of current sub-models evaluated in the multi-model ensemble risk projection tool with green lines representing 48 distinct combination of sub-models. Distinct paths through the ensemble are highlighted as black (NSCR2020), red (UNLV2017), purple (ORCRA2017), and green (RadRAT) lines.

The colored boxes along the bottom of the figure identify the major components of REID being represented with multiple sub-models and each line represents an ensemble REID estimate passing through a distinct combination of sub-models. NSCR2020 is currently considered NASA’s best estimate of individual sub-model implementation (black line) and is considered the “anchor model” of the ensemble. Additional distinct paths through the ensemble framework have been identified for comparison with the anchor model and full ensemble distribution as illustrated by the highlighted purple, red, and green lines.

The resulting probability density functions and cumulative probability functions formed by the individual ensemble members for the ~6-month lunar DRM are shown in Fig 7. Fig 7A shows the probability densities for the ensemble median, anchor model, and bounding ensemble members. For these analyses, the ensemble distribution assumes equal weighting of the 48 combinations (See discussion in Section V.III *Ensemble weighting*). The *R*_med_ and *R*_95%_ of the bounding cases are identified in Fig 7A as well as an indication of the sub-model path through the ensemble framework where sub-models differing from NSCR2020 are explicitly identified for each of the major components of uncertainty (Q, DDREF, ER, latency). Cumulative probabilities for the ensemble distribution (color contour), ensemble median, and anchor model are shown in Fig 7B. The ensemble median and anchor model fall near the middle of the path formed by the collection of individual members. The ensemble distribution was calculated from the individual ensemble member cumulative probabilities as follows. For a given percentile (*y*-axis of panel B), the inverse cumulative probability function of each ensemble member is evaluated. This creates a set of 48 discrete REID values from which a continuous cumulative distribution can be estimated.

**Fig 7.**
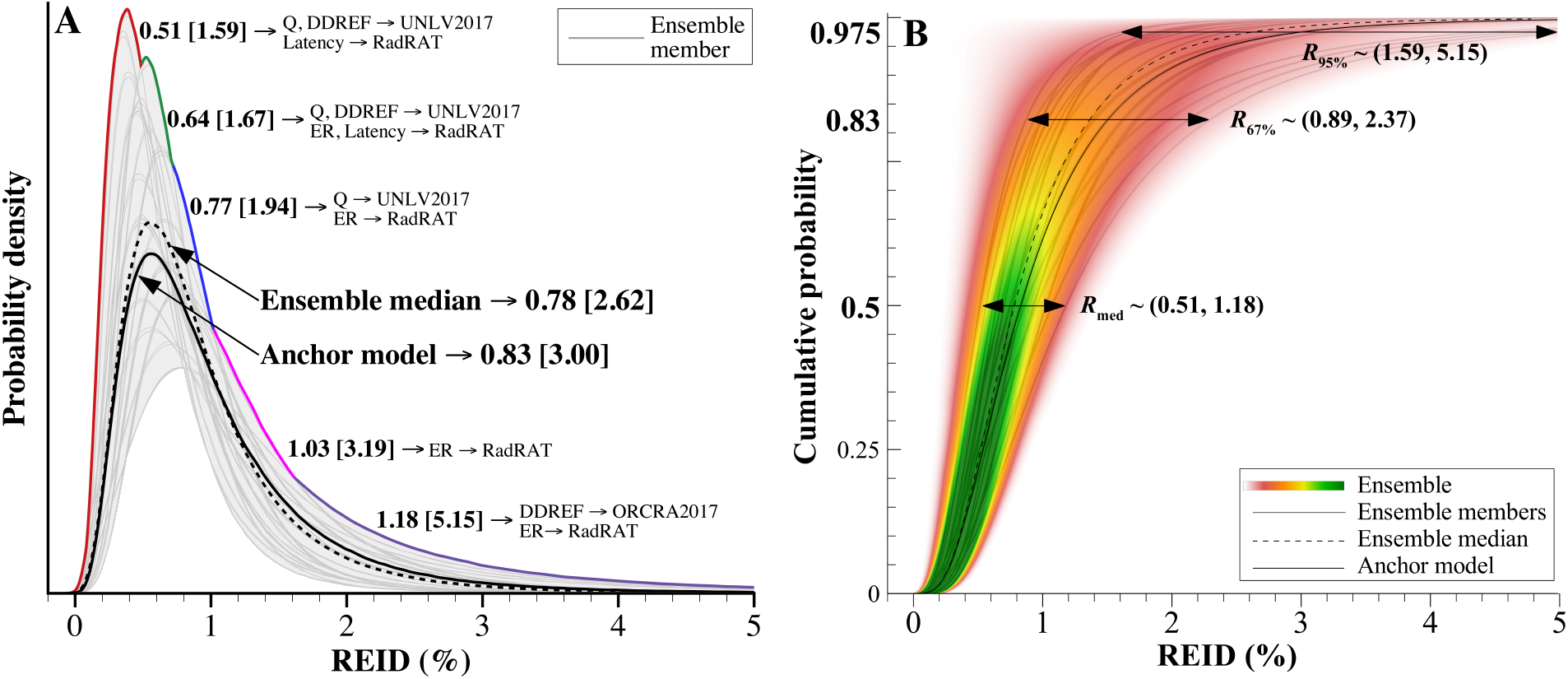
Ensemble risk of exposure induced death (REID) distribution for ~6 month lunar orbital design reference mission (DRM) with an effective dose of 185 mSv. Specifics of the DRM were selected such that the upper 95% CL REID value (*R*_95%_) aligns with the 3% PEL for direct comparison with NASA’s current limit. (A) Probability densities for the anchor model (solid line), ensemble median with equal sub-model weighting (dashed line), and ensemble extremes. Bounding ensemble member values are provided in the plot as *R*_med_ [*R*_95%_]. (B) Ensemble cumulative probabilities with an estimated *R*_med_ of 0.78%, *R*_67%_ of 1.40%, and *R*_95%_ of 2.62%. The contour heat map is obtained from a kernel density estimation of the ensemble members, where green is used to indicate increased member agreement relative to red.

Kernel density estimation (KDE) has been used in this initial phase of development to describe the continuous distribution with equal weight given to each ensemble member. The KDE bandwidth parameter was set as Silverman’s rule of thumb [Silverman 1986] with a scaling factor of 1.5 to ensure all ensemble members fell within the KDE distribution at each percentile. Generating these probability distributions over the range of percentiles from 0 to 1 yields a continuum of distributions shown as the contour plot in Fig 7B. We also obtain a median value at each percentile to form the ensemble median (dashed lines). Areas of dark green indicate close agreement amongst the ensemble members while areas of light red indicate increased dispersion. Note, the color contour indicates regions of consensus amongst ensemble members and should not be interpreted as the most probable REID value. Alternate methods for combining member and sub-model results into an ensemble distribution are discussed in Section V.III.

As discussed, the ensemble risk framework extends our understanding of uncertainties to include model-form contributions. Each of the single model ensemble members are able to characterize parameter uncertainties (Fig 7A) as the width of a given probability density function (i.e. fold-factor of *R*_95%_/*R*_med_ discussed in Section II.I). The bounding probability density functions highlight the broad range of *R*_95%_ and *R*_med_ values calculated by the relatively limited set of sub-models being considered. These variations are largely driven by quality and DDREF sub-models [Simonsen and Slaba 2020]. Model-form uncertainty is more easily seen as the width of the contour at a given cumulative probability (Fig 7B). For example, the range of ensemble member *R*_med_ values (found at a cumulative probability of 0.5) vary between 0.51% and 1.18%. The shaded green contour in this region suggests some level of consensus amongst the members relative to much greater dispersion at higher confidence levels. The range of *R*_95%_ values (found at a cumulative probability of 0.975) vary between 1.59% and 5.15%. While the lunar DRM mission duration of 174 days (~ 6 months) was specifically selected to align with the current limit (i.e., 3% at 95^th^ CL utilizing NSCR2020), under the same shielding and solar conditions the weighted-ensemble model projects a lunar orbital PMD of 199 days with individual ensemble member projections ranging between 101 and 328 days (for a 35 yr-old female never smoker). The ensemble results of Fig 7 provide visual evidence of additional uncertainty in REID calculations at the upper 95% CL that are not accounted for by any single model or by the current NASA PEL. Techniques to combine parameter and model-form uncertainties into a single scalable quantity are evolving and will necessarily implement advanced statistical methodologies [Hubin and Storvik 2019].

A qualitative comparison between the anchor model, ensemble members, and ensemble distribution clearly illustrates the spread of *R_med_* and *R*_95%_ values within the ensemble. While likelihood metrics have been used in other applications of ensemble forecasting to communicate forecast skill and/or degree of certainty of member models [Ray and Reich 2018; Kelly et al. 2019], there is a lack of statistically significant data for human space radiation health effects data to assess or score individual model prediction on actual outcome. However, application of similar methods may be used to statistically evaluate the level of agreement between models or deviation from the ensemble median. The Jensen-Shannon (JS) divergence [Schutze and Manning 1999] can be used to assess the overall level of agreement between two probability distributions and provides a simple metric for simultaneously comparing all of the distinct ensemble members to the ensemble median. The JS divergence between two probability density functions, *f* and *f*_0_, is calculated as

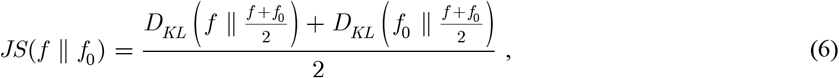

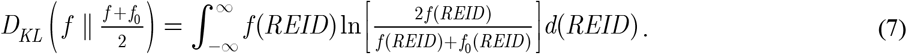

The quantity *D_KL_* is the Kullback-Leibler divergence [Kullback 1978], or relative entropy for two distributions. For identical distributions, the JS divergence is zero, while larger values of JS indicate growing differences between the distributions being compared. Here, the ensemble median is taken as *f*_0_, and the JS divergence is calculated for each of the ensemble members.

Fig 8 provides the JS divergence values for individual ensemble members compared with the ensemble median. The two latency models yield nearly identical REID distributions with JS divergence values exhibiting the same behavior. Therefore, only results incorporating the NSCR2020 latency model are shown in Fig 8. The path formed by NSCR2020 sub-model options (red text) produces a REID distribution that is closest to the ensemble median (smallest JS value). Also evident are the large JS divergence values associated with the UNLV2017 and ORCRA2017 DDREF models. This perceived divergence is not surprising and highlights the value of incorporating models based on different formalisms or assumptions. Additional independent models are needed to provide further insight into model-form uncertainties and strengthen ensemble projections. While these comparative measures confirm intuitively what we expect, an unbiased means to quantify the degree of relative agreement between ensemble member distributions will become increasingly important as future models employing different approaches are added.

**Fig 8.**
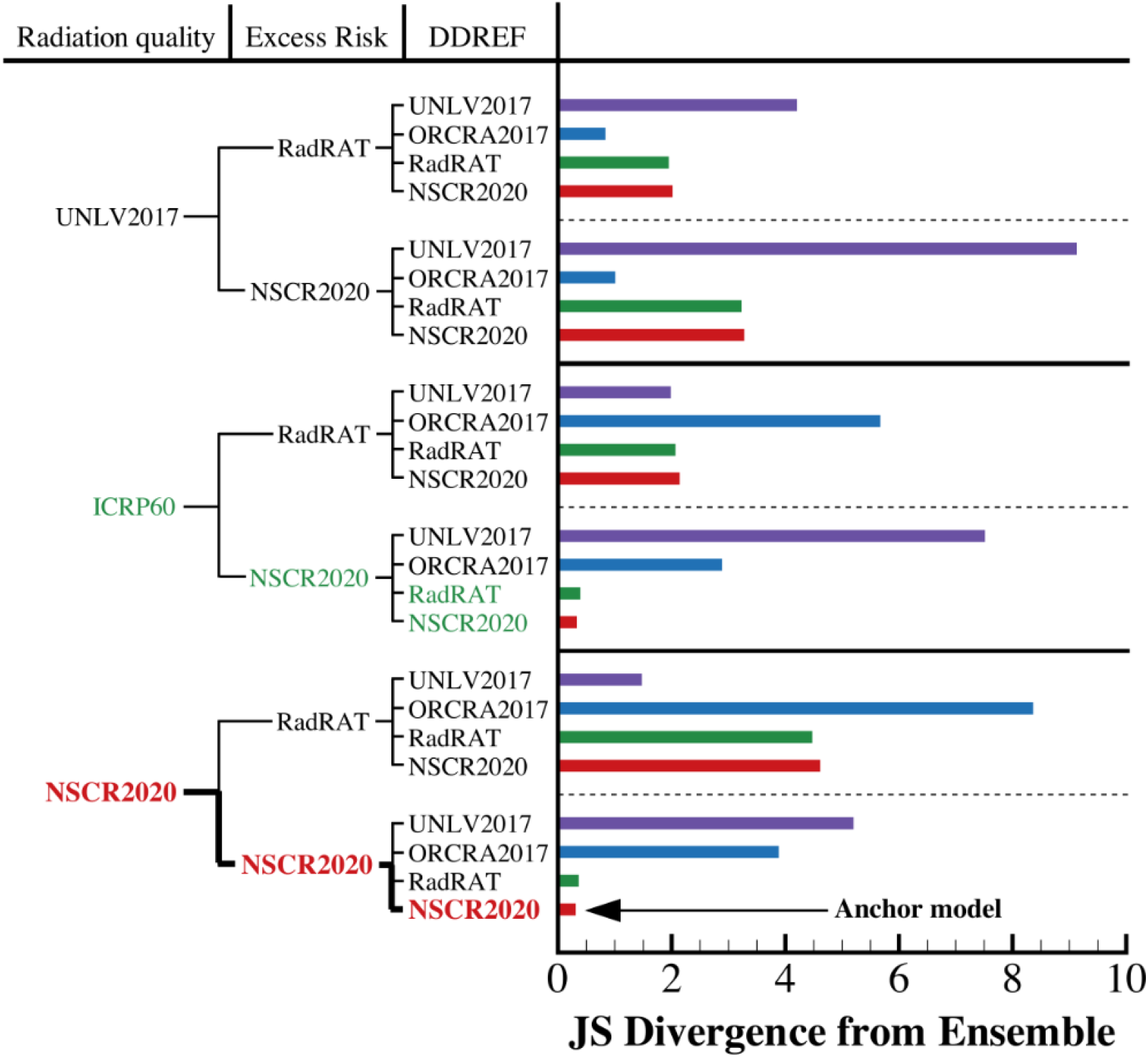
Comparison of distinct ensemble member REID distributions to the ensemble median using JS divergence as a measure of overall agreement. NSCR2020 (bottom row, bold red text) and two ICRP60 quality paths (green text) provide the closest agreement with the ensemble median.

### IV.II Utilizing ensemble methodologies to inform risk-based decisions

Ensemble-based risk projections offer a broader understanding of uncertainties which can support the focus of decision making toward regions with more stability and certainty and shift focus away from highly sensitive and uncertain prediction regions. Figs. 3 and 7 have shown that the *R*_95%_ value is itself uncertain and sensitive to underlying model assumptions. This suggests that a broader perspective of uncertainties in risk estimates may be needed to avoid a false sense of precision becoming embedded in the decision-making process for risk acceptance. For example in Fig 9, we show the KDE ensemble distributions obtained at the 50^th^, 83.3^rd^ (upper 67% CL) and 97.5^th^ (upper 95% CL) percentiles of Fig 7B for the lunar orbital DRM and now a 621-day Mars mission. The spread of ensemble member median values, represented by the green distributions, covers a relatively narrow REID interval. This level of agreement is expected since the models considered here are fundamentally similar and anchored to similar experimental and epidemiological data. Of greater interest is that model projections at the upper 67% CL become increasingly spread apart (orange distributions) covering a broader REID interval. Even greater divergence is seen in the distributions of upper 95% CL values (red distributions) where models are forced to increasingly extrapolate beyond limited biological data sets (See also Kappa discussion in Section II.I Challenges in space cancer risk projection). The width of each distribution can be interpreted as an initial measure of model-form uncertainty.

**Fig 9.**
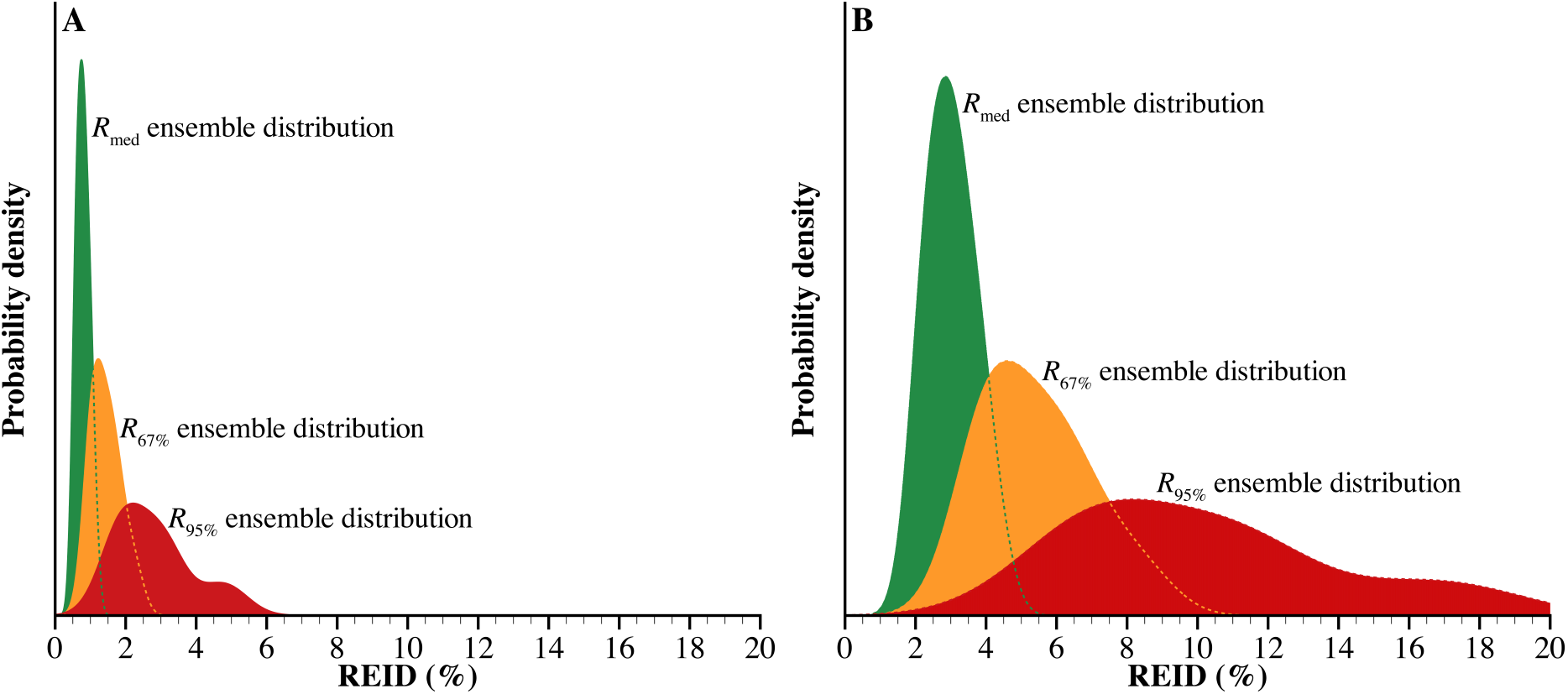
Ensemble distributions of the risk of radiation exposure induced death (REID) for the median, upper 67% CL, and upper 95% CL for exploration design reference missions (DRM). A) ~6 month lunar orbital DRM with an effective dose of 185 mSv and an estimated *R*_med_ of 0.78%, *R*_67%_ of 1.40%, and *R*_95%_ of 2.62%. B) Mars mission DRM with an effective dose of ~660 mSv and an estimated *R*_med_ of 2.97%, *R*_67%_ of 5.25%, and *R*_95%_ of 9.62%. Mars DRM assumes 621 days in transit with a short surface stay of 30 days for a 35 year-old female never-smoker behind 20 g/cm^2^ of aluminum shielding at solar minimum conditions.

## V. Discussion

Previous modeling efforts have focused on reducing uncertainty through refinements to the individual sub-models of NSCR2020 as a means to substantially increase permissible mission durations. However without major advancements in our scientific knowledge about the relationship between risk and dose, reducing uncertainties by several fold, as envisioned for Mars missions, is not achievable. Here, we shift our modeling focus to emphasize enhanced strategies for operational decision-making and mission planning when faced with these large uncertainties.

Ensemble forecast methodologies provide a means to systematically compare multiple predictive models, sub-models and uncertainty estimates, as well as influence the development of new models. Increased efforts to develop independent or alternate models with unique formalisms can improve the characterization of model-form uncertainties in regions where significant relevant data sets exist – such as near the median or at lower confidence intervals [Tebaldi and Knutti 2007]. However, additional independent models cannot significantly reduce uncertainties in regions that extrapolate beyond current observations (e.g., upper 95% CL) without a significant amount of new and relevant experimental and epidemiological data to inform development. Likewise, these new data can support the reduction of parameter uncertainties at larger confidence intervals. Additionally, models that account for the known uncertainties not currently represented in risk projections (e.g., gray boxes of Fig 5) will be needed to better inform risk-based decision making as mission durations and doses considerably increase beyond our current ISS experience base. Future Value of Information (VoI) analyses and Expected Value of Perfect Information (EVPI) methods can be used to guide research to understand the potential value of resolving uncertainty between model projections [Li et al. 2019].

### V.I Addressing model-form uncertainty

The ensemble framework developed here makes use of available sub-models to project radiogenic cancer risks. The current ensemble results do not yet incorporate a large number of independent sub-models implementing significant differences in fundamental assumptions or theories. For the ER models, the variation in REID was small compared to other factors, and is mainly attributed to differing age-dependencies in the parametric representations used by BEIR VII, UNSCEAR, and Preston and colleagues [Preston et al. 2007]. For the DDREF models, the variation shown is largely attributed to data selection in the UNLV2017 model and the use of ensemble methodologies that increased the likelihood of inverse dose-rate effects in the ORCRA2017 model. In the case of radiation quality, the UNLV2017 model is based on the same biological action cross section and initiation/promotion model as NSCR2020. However, the added coupling between DDREF and radiation quality in the HZE penumbra region of the track distinguishes the two models – with the assumption that dose rate effects are negligible in the track core - and leads to significant consequences on REID estimates. The radiation quality model comparisons provide an initial look at addressing model-form uncertainty and quantifying the impact of model assumptions on overall risk posture. With the current models largely based on the same data sets, these comparisons are especially beneficial in situations where one assumption may be just as valid as another but, as shown, can have large implications at the 95^th^ CL.

Inclusion of additional sub-models relying on fundamentally different approaches can further address model form uncertainty. Current models for ER rely mainly on LNT to extrapolate epidemiological data to low dose; however, alternate dose response models using multi-model inference techniques have been developed for specific tissues [Kaiser and Walsh 2013], and additional epidemiology data sets are being evaluated that can easily be implemented within this framework. Other models of radiation quality can be considered in future efforts, including the potential utilization of a radiation effects ratio (RER) in a narrow mission-relevant dose region [Shuryak et al. 2017], mixed-field quality factors, and coupled mixed-field/dose-rate multiplicative factors. Results from on-going research studies at the NSRL and the Colorado State low dose neutron facility [Simonsen et al. 2020, Borak et al. 2019] will supply a body of evidence to further inform model-form structure and the weighting (other than equal weighting) of various sub-model contributions accounting for mixed-field quality and dose-rate effects.

Additional model development and relevant radiobiology data sets will be needed to further quantify known uncertainties (model form error) not currently accounted for in risk prediction (gray boxes of Fig 5). For example, NSCR2020 assumes similarities in the spectrum and aggressiveness of tumors arising from high- versus low-LET irradiations while reported results using various animal models differ in outcome [Edmondson et al. 2020, Datta et al. 2013, Trani et al. 2010, Alpen et al. 1993]. Likewise, it is assumed that the biological response from a mixed-field irradiation (in particle type and energy), as would be found in space, can be determined by simple additivity or by directly adding responses from individual ions and energies. However, results for Harderian gland tumorigenesis [Siranart et al. 2016] and lung cancer tumorigenesis [Luitel et al. 2020] in murine models suggest that synergies between particles may exist and that more complex additivity models need to be considered. Emerging models for non-targeted effects (NTE), which have a significant influence on low-dose extrapolations expressed through RBEs, are being developed and have been shown to significantly impact radiation quality factors [Cucinotta and Cacao 2017, Matsuya et al. 2018]. Given the large cohorts and associated costs of low-dose studies, a general lack of animal data exists across high-risk tissues, doses, and particle types from which to resolve NTE model parameters [Chappell et al. 2020; Cucinotta et al. 2017]. Collectively, these examples imply that scaling procedures used to compute REID impose some level of uncertainty that is not currently accounted for in the present models.

### V.II Future ensemble members

While the current number of sub-models and distinct models may be limited, numerous development activities are underway that will provide additional data-sets and methods for inclusion (red boxes of Fig 5) given similar endpoints of REIC or REID. Ideally, model-form error can best be addressed by incorporating additional risk projection models that: 1) are independently developed (e.g., international models); 2) utilize different underlying epidemiology (e.g., Million Person Study, INWORKS); and/or 3) consider vastly different modeling approaches (e.g., biologically-based) and different modeling concepts as described above (e.g., non-targeted effects, RER vs. RBE, mixed-field quality factor).

Future ensemble modeling will compare results from ongoing epidemiological investigations conducted by the National Council on Radiation Protection (NCRP) including the “One Million U.S. Radiation Worker and Veteran Study [Boice, 2012, 2014; Bouville et al. 2015]” and the “*Evaluation of Sex-Specific Differences in Lung Cancer Radiation Risks and Recommendations for Use in Transfer Models (SC 1-27).*” These studies aim to reduce uncertainties in the projected REID and confidence interval by inclusion of a large US population of healthy workers who are more representative of the astronaut corps (e.g., similar with respect to health, ethnicity, and lifestyle factors) and consideration of protracted rather than acute exposures compared with the 1945 Japanese atomic bomb survivors. The Million Person Study (MPS) is a large-scale epidemiological investigation of one million U.S. radiation workers and atomic veterans (those exposed to ionizing radiation while present in the site of a nuclear explosion during active duty). The MPS population is 20 times larger than the adult Japanese study population [Ozasa et al. 2012] with individual exposures having cumulative doses >100 mSv at more space-relevant exposure rates [Bouville et al. 2015]. Inclusion of these data has the potential to reduce the uncertainties in the REID by removing the need to adjust for differences between a Japanese and a Western population and by minimizing or reducing the need for a DDREF adjustment [NCRP 2014]. The “*Sex-Specific Differences in Lung Cancer*” study will specifically evaluate the extent to which a sex difference exists across multiple available exposed populations and provide recommendations as to whether changes should be made in the sex-specific lung cancer risk coefficients used when transferring risks from one population to another. Thus, narrower probability distributions can presumably be achieved by removing the need to adjust for population differences (transfer function weighting), by utilizing additional epidemiological data in assigning sex-specific risk factors, and by minimizing or reducing the need for a DDREF adjustment.

Likewise, sub-models of excess risk, latency, and dose-rate extrapolation can include results from on-going international studies including those of the Inter Agency for Research on Cancer (IARC) and the ICRP. The IARC is coordinating epidemiology studies of international workers in the nuclear sector (INWORKS) to examine the risks of cancer and non-cancerous diseases linked to chronic ionizing radiation exposure at low doses and low dose-rates in cohorts of French, British, and American workers [Hamra *et al.* 2015]. The ICRP is currently reviewing available data on the estimation of risk at low doses in their Task Group 91, “*Radiation Risk Inference at Low-dose and Low-dose Rate Exposure for Radiological Protection Purposes*.” This report will make recommendations on alternative approaches of assessing the slope of dose response at high doses and then applying a DDREF reduction factor; or by inferring the risk coefficients at low doses using all available information and techniques of Bayesian analysis for estimating the best expert judgment.

In addition to the further development and handling of sub-models, available “full” risk models can be included in the ensemble such as those models under consideration by ESA and JAXA (given similar endpoints of REIC or REID) [Walsh et. al 2019, McKenna-Lawlor 2014]. For example, ESA is promoting the development of a space radiation risk model based on European-specific expertise in transport codes, radiobiological modeling, risk assessment, and uncertainty analysis for both cancer and non-cancer endpoints in support of exploratory class missions [Walsh et al. 2019]. Comparative analyses of models can support the ICRP’s evaluation on how to harmonize international models and dose limits [ICRP 2019]. These newly developed models may be included as distinct models within the ensemble for direct comparison to the NSCR2020 anchor model and weighted ensemble results. This is somewhat analogous to the National Hurricane Center’s ensemble of independent models to forecast the cone of hurricane tracks whereby predictions from two of its most well-known models, the U.S. National Weather Service’s Global Forecast System and the European Centre for Medium-Range Weather Forecasts, are distinctly identified.

An area of future work will focus on the development of sub-model/model entrance and exit criteria into the full ensemble forecast.

### V.III Ensemble weighting

A central issue in ensemble modeling is how to weight and combine projections to support decision making or action. Even a simple average of ensemble member predictions often produces a more skillful forecast than any individual ensemble member with the variation or spread of predictions providing a measure of uncertainty. Taking this approach a step farther, “corrected” consensus models assign different weights to each member model in an attempt to account for bias or systematic errors of individual members. Bayesian techniques used in weather forecasting have been successfully extended to epidemiological applications wherein multiple projections from a single parametric model are combined to compare the efficacy of various control actions on disease outbreaks [Lindström et al. 2015, Park et al. 2017]. Probabilistic projections of disease have also been scored using a log-likelihood (ignorance) score or mean log scores when comparing projected outbreak size to actual size [Kelly et al. 2019, Ray and Reich 2018].

Although in principle, the ideas of ensemble forecasting applied to weather and disease/immunization applications seem reasonable to extend to astronaut risk projection, there are limitations. Weather applications rely on a wealth of data to validate past performance of ensemble members, assign model weights, and set entrance and exit criteria within the ensemble forecast. Even with 20 years of International Space Station operational experience, there remains a statistically small number of individuals exposed to space radiation. Thus, REID cannot be directly measured or evaluated in the astronaut cohort, making it difficult to apply these same ideas in ensemble cancer risk projection. Additional animal data sets on dose-rate and quality effects, although limited, may support weighting of select sub-model members; however, the fact that many sub-models already rely on similar or identical data sets hinders a priori sub-model weight assignment. While the choice of equal weights as implemented here is subjective, it provides a reasonable starting point for software development.

Additional methods to improve the integration of ensemble member results are currently being considered. One method is a direct extension of the current formalism, wherein equal weighting of the paths taken within the code (Fig 6) can be replaced with subjectively assigned weights to account for perceived sub-model errors and bias. In this approach, individual sub-models can be assigned weights based on direct comparison to available data, subject matter expert opinion, and other considerations related to model maturity and reproducibility of results. However, the assignment of weights to individual sub-models will carry with it a host of questions which ultimately must be resolved with potentially an unwanted degree of subjective decision-making. A second method takes a more statistical approach by considering the region of overlap between multiple predictive models from which to infer uncertainty and determine weights objectively. Bayesian model averaging or combination is a possible avenue by which ensemble member weights may actually be inferred, instead of specified [Fragoso et al. 2018]. A complicating feature of these methods, however, is that the underlying members making up the ensemble must be compared against some true, or measured, value(s). Other methods may consider using log-likelihood or mean log scoring of probabilistic projections to evaluate likely agreement or deviation from NASA’s operational model. Analyses will necessarily focus on evaluating the extent to which models agree and on understanding the underlying assumptions driving large uncertainties where models deviate. Alternate approaches for ensemble member weighting and combinatorial methods will be a priority of future research efforts.

### V.IV Application of ensemble methods for risk reporting and potential PEL updates

NASA’s current PEL has been in place since 2003 [NASA 2014] and was deemed both “reasonable” and “achievable” given the corps of astronauts and the types of LEO missions that NASA was performing. In 2014, a supplementary review found the application of the 95% CL to the 3% REID for cancer prudent and appropriate for LEO missions given the currently accounted for uncertainties in knowledge about the biological effects of space radiation (Table 1) and the potential implications of additional uncertainties not currently accounted for in NASA’s risk model [NCRP 2014]. With new missions being planned beyond LEO, NASA OCHMO, per established processes [NASA 2016], is reviewing the applicability of the current radiation health standard (https://www.nationalacademies.org/our-work/assessment-of-strategies-for-managing-cancer-risks-associated-with-radiation-exposure-during-crewed-space-missions). Additionally, for consistency in risk communication with other spaceflight health risks [NASA 2013], NASA is considering radiation risk reporting of likelihood using a measure of central tendency (e.g., *R*_med_) with the confidence interval presented a measure of statistical plausibility to ensure clinical relevance. This is in contrast to risk communication solely at the 95^th^ CL - which may actually provide a false sense of certainty given that the REID at such a high CL is a highly sensitive quantity and heavily influenced by subjective assumptions and incomplete biological data as illustrated in these analyses.

Here we have more comprehensively characterized radiogenic cancer risk through ensemble methodologies based upon available published models and epidemiology. Parameter uncertainties and now model-form uncertainties are considered to support decision-making when faced with the large uncertainties surrounding the biological effects of space radiation. Several important conclusions can be drawn: 1) as a measure of ensemble model central tendency, the distribution of medians (*R*_med_) is narrow (Fig 9) with underlying models in relative agreement (with the caveat that they are largely based on similar assumptions and data); 2) the distribution of upper 67% CL values (*R*_67%_) includes conservatism to account for uncertainties ascertained by limited experimental and epidemiological data with only moderate dependence on underlying model assumptions; and 3) large model-form uncertainties exist where the current PEL is defined (*R*_95%_) and are largely driven by subjective assumptions lacking robust experimental and epidemiological support (Fig 9). Applications of these results impacting risk reporting and PEL definition are offered as *illustrative* examples.

#### V.IV.I Risk Reporting

For risk reporting to flight surgeons and crew, the ensemble median with a range of values defined at the upper 67% CL and upper 95% CL may better communicate our current state of knowledge and known uncertainties. For the 6-month lunar orbital DRM considered throughout this work, the ensemble-based risk projection (for a 35-year-old female) would be reported as a median REID of 0.78%±0.21% with an informed upper bound of 1.40%±0.47% and a conservative upper bound of 2.62%±1.18. Here, “informed” and “conservative” upper bounds refer to the upper 67% CL and upper 95% CL, respectively. In addition, to communicate variation within the *R*_med_, *R*_67%_, and *R*_95%_ distributions (green, orange, red distributions, respectively of Fig 9A), an additional measure of uncertainty has been added to reflect the variation away from the medians of each of these distributions set at the 67% confidence interval. For the 621-day Mars mission DRM (for a 35-year-old female), the risk projection would be reported as a median REID of 2.97%±0.79% with an informed upper bound of 5.25%±1.72% and a conservative upper bound of 9.62%±4.06% (with values informed from in Fig 9B). The understanding of these uncertainties is required in communicating risk to crew for informed consent and personal clinical management and equally to those responsible for decisions in allowing (or not) higher risk to crew to meet exploration mission objectives. Given the large uncertainties in the biological effects of space radiation, risk reporting solely at the median does not include sufficient information to ensure adequate protection of crew long-term health.

#### V.IV.II Permissible Exposure Limits

In the near term, the ensemble risk projection offers additional information to support potential updates to NASA’s PEL - in spite of the limited number of sub-models available. *As a notional example*, a revised PEL that maintains nearly the same level of risk tolerance as the current PEL (*R*_med_ on the order of 1%) could be considered such that: “planned career exposures do not exceed a 2% REID for cancer mortality evaluated at a 67% confidence level to limit the cumulative effective dose (in units of Sievert) received by an astronaut throughout his or her career.” In this example, the confidence level has been adjusted downward from 95% to 67% because of the large ensemble model divergence at the upper 95% CL (Fig. 9) and its sensitivity to subjective assumptions (Fig. 3). In both cases, the ensemble prediction of a 3% REID at the 95^th^ CL and a 2% REID at the 67^th^% CL provide an estimated *R*_med_ on the order of 1%; however the 2% REID at 67^th^ CL is a more stable quantity amenable to operational implementation. Similarly, higher (or lower) risk acceptance (*R*_med_) at other CLs (presumably less than the 95^th^) can easily be evaluated using these methodologies to inform PEL updates. While implementation of an ensemble framework with the addition of model-form uncertainties may seem to further complicate PEL definition, in an operational setting it is quite the opposite whereby the PEL can now be defined in a more stable region of the probability distribution function where parameter and model-form uncertainties are narrower. Similar to weather and disease prediction,

Consistent with other spacefaring nations who implement dose-based limits, this methodology can reliably inform a dose-based PEL system based on a risk posture that NASA deems acceptable. In the current example (with *R*_med_ equal to approximately 1%), a 35-year-old female’s planned career exposure would not exceed a cumulative effective dose of 185 mSv to limit cancer mortality to < 2% REID at the 67^th^ CL. This maintains a relationship between risk and dose such that as major advancements in scientific knowledge and our understanding of uncertainties evolve, the physical quantity of dose can be modified in standard updates. This is in contrast to above where %REID remains the defined PEL quantity and exposure (mSv) is operationally controlled through risk projections and PMDs. Defining the CL between >50% to <95% will allow for an increased exposure (mSv) limit if substantial reductions in uncertainty are realized with the greatest impact at the highest set CL. Thus, a balance would need to be maintained in this region – that is, not setting the CL artificially low such that advancements in knowledge (uncertainty reduction) barely influence limits or too great where subjective decisions, not anchored by sufficient empirical evidence, dominate. Ensemble analyses such as these can provide insight to help define an appropriate CL to adequately account for both parameter and model-form uncertainties.

In reviewing the applicability of the current health standard and available clinical evidence base, the specific numerical values (%REID, CL, and/or effective dose) will need to be specified to reflect NASA’s acceptable risk tolerance for missions beyond LEO. In these examples, central tendency is reported, and the PEL is anchored to a region with greater certainty based on existing evidence. Additional reporting at the upper 95^th^ captures an indication of uncertainty levels due to incomplete biological knowledge consistent with NCRP recommendations.

## VI. Summary

A new approach to NASA risk modeling has been developed to extend the current NASA probabilistic cancer risk model to a multi-model ensemble framework capable of assessing sub-model parameter uncertainty as well as model form uncertainty associated with differing theoretical or empirical formalisms. This hybrid PP/MM approach has been specifically applied to the dominant terms in the astronaut risk projection sub-models, including the radiation quality factor, DDREF, latency function, and excess risk functions, with each sub-model containing its own description of parameter uncertainty. The sensitivity of subjective model assumptions contributing to uncertainty at the 95% CL can be readily evaluated within the ensemble framework to inform PELs, acceptable permissible mission durations, and risk posture for crew. Given the models considered, NSCR2020 was found to be a reasonable estimate of cancer risk projection with a median and 95% CL close to the ‘equal-weighted’ ensemble model prediction. This general agreement is expected and is largely due to the underlying sub-models implementing similar approaches and reliance on limited epidemiology and relevant experimental data sets. However, in assessing parameter uncertainty and now model-form uncertainty jointly, a broad range of values exist in the probability distributions at the upper 95% CL where the NASA PEL is defined. This is largely driven by model extrapolations beyond our state of knowledge. Thus, crew permissible mission durations are being defined in a very dynamic portion of the REID probability distribution while a comparison of ensemble distributions at lower CL’s provides comparatively narrower estimates where multiple models are in better agreement.

Emerging data sets from the NSRL are necessary to advance sub-model development and to support the future weighting of ensemble members based on experimental evidence [Simonsen and Slaba 2020]. Inclusion of additional risk projection models that are independently developed (e.g. international models), consider vastly different modeling approaches (e.g. biologically-based), utilize different underlying epidemiology (i.e. US million person data), and/or different modeling concepts (e.g. non-targeted effects, RER vs. RBE, mixed-field quality factor) can further improve our understanding of uncertainty in the ensemble forecast. Future work will incorporate new models and data sets which meet model entrance criteria and incorporate the appropriate rigorous statistical methods to combine and/or weight multiple risk projections.

In the near term, utilizing the methodologies described here can provide a more complete picture of our state of knowledge and associated uncertainties the risk landscape should PEL changes be pursued for NASA’s future Artemis missions. In the longer term, ensemble modeling can provide crew, flight surgeons, and policy decision-makers additional information to reach decisions on risk acceptance for long-duration exploration missions such as human missions to Mars. This will be particularly important if current health and medical standards cannot be met or the level of knowledge does not permit a standard to be developed. These efforts support a rigorous process to assure that crew are fully informed about risks and unknowns as described by the Institute of Medicine’s’ report, *Health Standards for Long Duration and Exploration Spaceflight: Ethics Principles, Responsibilities, and Decision Framework*” [IOM 2014].

## VII. Acknowledgements

This work was performed by the Multi-Model Ensemble Risk Assessment (MERA) project at NASA Langley Research Center and is supported by the Human Research Program under the Human Exploration and Operations Mission Directorate at NASA.

## Footnotes

1 Previous analyses have assessed the modification of the free-space GCR environment through both complex spacecraft (such as the International Space Station) and simplified geometries to quantify the variability of the induced tissue field within critical body tissues. A simplified spherical shield of 20 g/cm^2^ aluminum provides a reasonable estimate of typical spacecraft shielding with a variation in dose (Gy) and dose equivalent (Sv) across all major radiosensitive tissues and geometries found to be ±3% and ±16%, respectively [Slaba et al. 2016]. Similar conclusions are reached from assessments of the modified GCR spectrum in terms of flux versus linear energy transfer (LET) [Slaba et al. 2016].

2 Linear energy transfer (LET) is defined as the energy lost per unit path length and is usually expressed in units of keV/μm.

## References

Adamczyk AM, Norman RB, Sriprisan SI, Townsend LW, Norbury JW, Blattnig SR, Slaba TC, NUCFRG3: Light ion improvements to the nuclear fragmentation model. Nucl. Instr. Meth. Phys. Res. A 678; 2012. pp. 21–32.

Adriani O, et al., PAMELA measurements of cosmic-ray proton and helium spectra. Science 332; 2011. pp. 69–72.

Adriani O, et al., Time dependence of the proton flux measured by PAMELA during the 2006 July – 2009 December solar minimum. Astrophys. J. 765; 2013. pp. 1–8.

Adriani O, et al., Ten years of PAMELA in space. Rivista del Nuovo Cimento 40; 2017. pp. 473–522.

Aguilar M, et al., Precision measurement of the helium flux in primary cosmic rays of rigidities 1.9 GV to 3 TV with the Alpha Magnetic Spectrometer on the International Space Station. Phys. Rev. Lett. 115; 2015a. 211101.

Aguilar M, et al., Precision measurement of the proton flux in primary cosmic rays from rigidity 1 GV to 1.8 TV with the Alpha Magnetic Spectrometer on the International Space Station. Phys. Rev. Lett. 114; 2015b. 171103.

Aguilar M, et al., Observation of the identical rigidity dependence of He, C, and O cosmic rays at high rigidities by the Alpha Magnetic Spectrometer on the International Space Station. Phys. Rev. Lett. 119; 2017. 251101.

Aguilar M, et al., Observation of new properties of secondary cosmic rays lithium, beryllium, and boron by the Alpha Magnetic Spectrometer on the International Space Station. Phys. Rev. Lett. 120; 2018a. 021101.

Aguilar M et al., Precision measurement of cosmic-ray nitrogen and its primary and secondary components with the Alpha Magnetic Spectrometer on the International Space Station. Phys. Rev. Lett. 121; 2018b. 051103.

Aguilar M, et al., Observation of fine time structures in the cosmic proton and helium fluxes with the alpha Magnetic Spectrometer on the International Space Station. Phys. Rev. Lett. 121; 2018c. 051101.

Alpen EL, Powers-Risius P, Curtis SB, DeGuzman R. Tumorigenic potential of high-Z, high-LET charged particle radiations. Rad. Res. 88, 1993. pp. 132–142.

Arias E. United States Life Tables, 2011. National Vital Statistics Reports, Vol. 64, No. 11, 2015.

Bahadori AA, Sato T, Slaba TC, Shavers MR, Semones EJ, Van Baalen M, Bolch WE, A comparative study of space radiation organ doses and associated cancer risk using PHITS and HZETRN. Phys. Med. Biol 58; 2013. pp. 7183–7207.

Barcellos-Hoff MH, Mao JH. HZE radiation non-targeted effects on the microenvironment that mediate mammary carcinogenesis. Front. Oncol. 11, 2016. [https://doi.org/10.3389/fonc.2016.00057]

Bennett J, Little MP, Richardson S. Flexible dose-response models for Japanese atomic bomb survivor data: Bayesian estimation and prediction of cancer risk. Radiat. Environ Biophys. 43, 2004. pp. 233–245. [doi:10.1007/s00411-004-0258-3]

Billings MP, Yucker WR, The computerized anatomical man (CAM) model. Summary Final Report, MDC-G4655, McDonnell Douglas Company; 1973.

Boice JD JR. A study of one million U.S. radiation workers and veterans. A new National Council on Radiation Protection initiative. Health Phys. News, November 2012. pp. 7–10. [https://ncrponline.org/wp-content/themes/ncrp/PDFs/BOICE-HPnews/Nov-2012_Million_Worker.pdf]

Boice JD JR, Cohen SS, Mumma MT, Ellis ED, Cragle DL, Eckerman KF, Wallace PW, Chadda B, Sonderman JS, Wiggs LD, Richter BS, Leggett RW. Mortality among mound workers exposed to polonium-210 and other sources of radiation, 1944-1979. Rad. Res. 181, 2014. pp. 208–228.

Borak TB, Krumland N, Heilbrom LH, Weil MM (2019). Design and dosimetry of a facility to study health effects following exposures to fission neutrons at low dose rates for long durations. Int. J. Rad. Bio.; 2019. [doi:10.1080/09553002.2019.1688884]

Bouville A, Toohey RE, Boice JD Jr, Beck HL, Dauer LT, Eckerman KF, Hagemeyer D, Leggett RW, Mumma MT, Napier B, Pryor KH, Rosenstein M, Schauer DA, Sherbini S, Stram DO, Thompson JL, Till JE, Yoder C, Zeitlin C. Dose reconstruction for the million worker study: status and guidelines. Health Phys. 108, 2015. pp. 206–220.

Chappell LJ, Elgart SR, Milder CM, Semones EJ. Assessing nonlinearity in harderian gland tumor induction using three combined HZE-irradiated mouse datasets. Rad. Res. 194, 2020. pp. 38–51. [doi:10.1667/RR15539.1]

Clette F, Cliver EW, Lefèvre L, Svalgaard L, Vaquero JM, Leibacher JW, 2016: Preface to topical issue: recalibration of the sunspot number, Solar Phys. 291: 2479–2486; 2016.

Cucinotta FA, Kim MY, Ren L. Evaluating shielding effectiveness for reducing space radiation cancer risks. Rad. Meas. 41, 2006. pp.1173–1185.

Cucinotta FA, Kim MY, Chappell LJ. Space radiation cancer risk projections and uncertainties – 2012. NASA TP 2013-207375, 2013.

Cucinotta, FA. A new approach to reduce uncertainties in space radiation cancer risk predictions. PloS one 10, 2015. e0120717. [doi:10.1371/journal.pone.0120717]

Cucinotta FA, Alp M, Rowedder B, Kim MY. Safe days in space with acceptable uncertainty from space radiation exposure. Life Sci. Space Res. 5, 2015. pp. 31–38.

Cucinotta FA, New Estimates of radiation risks are favorable for Mars exploration however major scientific questions remain unanswered. FISO Colloquium, 2016. http://fiso.spiritastro.net/telecon16-18/Cucinotta_7-13-16/

Cucinotta FA, Cacao E. Non-targeted effects models predict significantly higher Mars mission cancer risk than targeted effects models. Scientific Reports 7, 2017. pp. 1832.

Cucinotta FA, To K, Cacao EE. Predictions of space radiation fatality risk for exploration missions. Life Sci. Space Res. 13, 2017. pp. 1–11.

Cucinotta FA. Radiation health risks for a Mars mission. Mars Sustainability Workshop, Buzz Aldrin Research Institute, Kennedy Space Center, Florida, 2018.

Cucinotta FA, Cacao EE, Kim MY, Saganti PB. Benchmarking risk predictions and uncertainties in the NSCR model of GCR cancer risks with revised low LET risk coefficients. Life Sci. Space Res. 27, 2020a. pp. 64–73.

Cucinotta FA, Cacao EE, Saganti PB. NASA space cancer risk (NSCR) model 2020. COSPAR 2020. 2020b.

Datta K, Suman S, Kallakury BV, Fornace AJ. Heavy ion radiation exposure triggered higher intestinal tumor frequency and greater β-catenin activation than γ radiation in APC Min/+ mice. PLoS One 8, 2013. e59295.

Edmondson EF, Gatti DM, Ray FA, Garcia EL, Fallgren CM, Kamstock DA, Weil MM. Genomic mapping in outbred mice reveals overlap in genetic susceptibility for HZE ion– and γ-ray–induced tumors. Sci. Adv. 6, 2020. eaax5940.

Fragoso TM, Bertoli W, Louzasa F. Bayesian model averaging: a systematic review and conception classification. Int. Stat. Rev. 86, 2018. pp. 1–28.

Fritsch JM, Hilliker J, Ross J, Vislocky RL. Model consensus. Weather Forecasting 15, 2000. pp. 571–582. [doi: 10.1175/1520-0434(2000)015<0571:MC>2.0.CO;2]

de Gonzalez AB, Apostoaei AI, Veiga LHS, Rajaraman P, Thomas BA, Hoffman FO, Gilbert E, Land C. RadRAT: a radiation risk assessment tool for lifetime cancer risk projection. J. Radiol. Prot. 32, 2012. pp. 205–222.

Guerra JA, Murray SA, Bloomfield DS, Gallagher PT. Ensemble forecasting of major solar flares: methods for combining models. Journal of Space Weather and Space Climate 10, 2020. [doi: 10.1051/swsc/2020042]

Hamra GB, Richardson DB, Cardis E, Daniels RD, Gillies M, O’Hagan JA, Haylock R, Laurier D, Leuraud K, Moissonnier M, Schubauer-Berigan M, Thierry-Chef I, Kesminiene A. Cohort profile: the international nuclear workers study (INWORKS). Int. J. Epidemiol 2015. [doi:10.1093/ije/dyv122]

Hoel DG. Comments on the DDREF estimate of the BEIR VII committee. Health Phys. 108, 2015. pp. 351–356.

Hubin A, Storvik G. Combining model and parameter uncertainty in Bayesian neural networks. Mathematics, Computer Science ArXiv, 2019. [https://arxiv.org/abs/1903.07594]

ICRP, International Commission on Radiological Protection. Recommendations of the International Commission on Radiological Protection. ICRP Publication 60. Pergamon Press, 1991.

ICRP, International Commission on Radiological Protection. Basic Anatomical and Physiological Data for Use in Radiobiological Protection: Reference Values. ICRP Publication 89, Pergamon Press, 2001.

ICRP, International Commission on Radiological Protection. The 2007 Recommendations of the International Commission on Radiological Protection. ICRP Publication 103. Pergamon Press, 2007.

ICRP, International Commission on Radiological Protection. Terms of Reference for ICRP Task Group 115: “Risk and Dose Assessment for Radiological Protection of Astronaut;” approved by the ICRP Main Commission on 20 May 2019.

IOM, Institute of Medicine. “Health Standards for Long Duration and Exploration Spaceflight: Ethics Principles, Responsibilities, and Decision Framework.” Committee on Aerospace Medicine and Medicine in Extreme Environments. Washington DC, National Academies Press (US), 2014. [http://www.nap.edu/catalog.php?record_id=18576]

Kaiser JC, Jacob P, Meckbach R, Cullings HM. Breast cancer risk in atomic bomb survivors from multi-model inference with incidence data 1958–1998. Radiat. Environ. Biophys. 51, 2012. pp. 1–14. [doi:10.1007/s00411-011-0387-4]

Kaiser JC, Walsh L. Independent analysis of the radiation risk for leukaemia in children and adults with mortality data (1950–2003) of Japanese A-bomb survivors. Radiat. Environ. Biophys. 52, 2013. pp. 17–27. [doi:10.1007/s00411-012-0437-6]

Katz R, Ackerson B, Homayoonfar M, Sharma SC, Inactivation of cells by heavy ion bombardment. Rad. Res. 47, 1971. pp. 402–425.

Kelly JD, Worden L, Wannier SR, Hoff NA, Mukadi P, Sinai C, et al. Projections of Ebola outbreak size and duration with and without vaccine use in Équateur, Democratic Republic of Congo, as of May 27, 2018. PLoS ONE 14, 2019. e0213190. [https://doi.org/10.1371/journal.pone.0213190]

Kocher DC, Apostoaei AI, Hoffman FO, Trabalka JR. Probability distribution of dose and dose-rate effectiveness factor for use in estimating risks of solid cancers from exposure to low-LET radiation. Health Phys. 114, 2018. pp. 602–622.

Kotaro O, Shimizu Y, Suyama A, Kasagi F, Soda M, Grant EJ, Sakata R, Sugiyama H, Kodama K. Studies of the mortality of atomic bomb survivors, Report 14, 1950-2003: an overview of cancer and non-cancer diseases. Rad. Res. 177, 2012. pp. 229–243.

Kramer R, Vieira JW, Khoury HJ, Lima FRA, Fuelle D, All about MAX: a male adult voxel phantom for Monte Carlo calculations in radiation protection dosimetry. Phys. Med. Bio. 48; 2003. pp. 1239–1262.

Kramer R, Vieira JW, Khoury HJ, Lima FRA, Loureiro ECM, Lima VJM, Hoff G, All about FAX: a female adult voxel phantom for Monte Carlo calculations in radiation protection dosimetry. Phys. Med. Bio. 49; 2004. pp. 5203–5216.

Kullback S. Information Theory and Statistics, John Wiley & Sons. Republished by Dover Publications in 1968, reprinted in 1978: ISBN 0-8446-5625-9.

Li SL, Ferrari MJ, Bjørnstad ON, Runge MC, Fonnesbeck CJ, Tildesley MJ, et al. Concurrent assessment of epidemiological and operational uncertainties for optimal outbreak control: Ebola as a case study. Proceedings. Biological Sciences 286, 2019. [doi:10.1098/rspb.2019.0774]

Lindström T, Tildesley M, Webb C. A Bayesian ensemble approach for epidemiological projections. PLoS Comput. Biol. 11, 2015. e1004187. [https://doi.org/10.1371/journal.pcbi.1004187]

Little MP, Hoel DG, Molitor J, Boice JD, Wakeford R, Muirhead CR. New models for evaluation of radiation-induced lifetime cancer risk and its uncertainty employed in the UNSCEAR 2006 report. Rad. Res. 169, 2008. pp. 660–676.

Luitel K, Kim SB, Barrona S, Richardsonb JA, Shay JW. Lung cancer progression using fast switching multiple ion beam radiation and countermeasure prevention. Life Sci. Space Res. 24, 2020. pp 108–115. [https://doi.org/10.1016/j.lssr.2019.07.011]

Martucci M, et al., Proton fluxes measured by the PAMELA experiment from the minimum to the maximum solar activity for solar cycle 24. Astrophys. J. Lett. 854; 2018.

Stone EC, Frandsen AM, Mewaldt RA, Christian ER, Margolies D, Ormes JF, Snow F, The advanced composition explorer, Space Sci. Rev. 86; 1998. pp. 1–22.

Matsuya Y, Sasaki K, Yoshii Y, Okuyama G, Date H. Integrated modeling of cell responses after irradiation for DNA targeted effects and non-targeted effects. Scientific Reports 8, 2018, 4849.

Matthia D, Ehresmann B, Lohf H, Kohler J, Zeitlin C, Appel J, Sato T, Slaba TC, Martin C, Berger T, Boehm E, Boettcher S, Brinza DE, Burmeister S, Guo J, Hassler DM, Posner A, Rafkin SCR, Reitz G, Wilson JW, Wimmer-Schweingruber RF, The Martian surface radiation environment – a comparison of models and MSL/RAD measurements. J. Space W. Space Clim. 6; 2016. pp. A13.

Matthia D, Hassler DM, de Wet W, Ehresmann B, Firan A, Flores-McLaughlin J, Guo J, Heilbronn LH, Lee K, Ratliff H, Rios RR, Slaba TC, Smith M, Stoffle NN, Townsend LW, Berger T, Reitz G, Wimmer-Schweingruber RF, Zeitlin C, The radiation environment on the surface of Mars – summary of model calculations and comparison to RAD data. Life Sci. Space Res. 14; 2017. pp. 18–28.

McKenna-Lawlor S. Feasibility study of astronaut standardized career dose limits in LEO and the outlook for BLEO. Acta Astronautica 104, 2014. pp. 565–573.

Mertens CJ, Slaba TC, Hu S. Active dosimeter-based estimate of astronaut acute radiation risk for real-time solar energetic particle events. Space Weather 16, 2018. pp. 1291–1316.

Mertens CJ, Slaba TC. Characterization of solar energetic particle radiation dose to astronaut crew on deep space exploration missions. Space Weather 14, 2020. pp. 1650–1658.

NA/NRC, National Academies/National Research Council. Radiation Hazards to Crews on Interplanetary Missions: Biological Issues and Research Strategies. National Academies Press, Washington DC, 1996.

NA/NRC, National Academies/National Research Council. A Strategy for Research in Space Biology and Medicine in the New Century. National Academies Press, Washington DC, 1998.

NASA, National Aeronautics and Space Act. Public Law 85–568 (July 29), 72 Stat. 426, 1958. [http://history.nasa.gov/spaceact.html] (Accessed October 1, 2020) (National Aeronautics and Space Administration, Washington DC).

NASA, Human Research Program Integrated Research Plan, HRP-47065 Rev D, 2012. [http://www.nasa.gov/sites/default/files/651214main_Human_Research_Program_Integrated_Research_Plan_RevD.pdf] (Accessed October 1, 2020) (National Aeronautics and Space Administration, Houston).

NASA, Human Research Program Evidence, 2013. [http://humanresearchroadmap.nasa.gov/evidence] (Accessed October 1, 2020) (National Aeronautics and Space Administration, Houston).

NASA, NASA Space Flight Human System Standard. NASA STD 3001, Vol I; 2014.

NASA, Health and Medical Requirements for Human Space Exploration. NASA Procedural Requirements 8900.1B, 2016.

NASA, Artemis Plan: NASA’s Lunar Exploration Program Overview. National Aeronautics and Space Administration, Washington DC, Sept. 21, 2020. [https://www.nasa.gov/sites/default/files/atoms/files/artemis_plan-20200921.pdf] (Accessed October 1, 2020)

NCRP, National Council on Radiation Protection and Measurements. Guidance on Radiation Received in Space Activities. NCRP Report No. 98, Bethesda MD, 1989.

NCRP, National Council on Radiation Protection and Measurements. Recommendations of Dose Limits for Low Earth Orbit. NCRP Report 132, Bethesda MD, 2000.

NCRP, National Council on Radiation Protection and Measurements. Radiation Protection for Space Activities: Supplement to previous recommendations. NCRP Commentary 23, Bethesda MD, 2014.

NRC, National Research Council. Health risks from exposure to low levels of ionizing radiation. BEIR VII Phase 2 report. National Academies Press, 2006.

NRC, National Research Council. Committee for Evaluation of Space Radiation Cancer Risk Model, Technical evaluation of the NASA model for cancer risk to astronauts due to space radiation. National Academy of Sciences Press, Washington DC, 2012.

Norbury JW, Slaba TC. Space radiation accelerator experiments – the role of neutrons and light ions. Life Sci. Space Res. 3, 2014. pp. 90–94.

Norbury JN, Double-differential fragmentation (DDFRG) models for proton and light ion production in high energy nuclear collisions valid for both small and large angles. NASA TP 2020-5001740; 2020.

O’Neill PM, Foster CC, Kim MY, Badhwar-O’Neill 2011 galactic cosmic ray flux model description. NASA TP 2013-217376; 2013.

Ozasa K, Shimizu Y, Suyama A, Kasagi F, Soda M, Grant EJ, Sakata R, Sugiyama H, Kodama K. Studies of the mortality of atomic bomb survivors, Report 14, 1950-2003: An overview of cancer and non-cancer diseases. Rad. Res. 177, 2012. pp. 229–243.

Park J, Goldstein J, Haran M, Ferrari M. An ensemble approach to predicting the impact of vaccination on rotavirus disease in Niger. Vaccine 35, 2017. pp. 5835–5841. [https://doi.org/10.1016/j.vaccine.2017.09.020]

Pierce DA, Vaeth M. The shape of the cancer mortality dose-response curve for the A-bomb survivors. Rad. Res. 126, 1991. pp. 36–42.

Preston DL, Ron E, Tokuoka S, Funamoto S, Nisha N, Soda M, Mabuchi K, Kodama K. Solid cancer incidence in atomic bomb survivors: 1958-1998. Rad. Res. 168, 2007. pp. 1–64.

Ray EL, Reich NG. Prediction of infectious disease epidemics via weighted density ensembles. PLoS Comput. Biol. 14, 2018. e1005910. [https://doi.org/10.1371/journal.pcbi.1005910]

Schöllnberger H, Eidemüller M, Cullings HM, Simonetto C, Neff F, Kaiser JC. Dose-responses for mortality from cerebrovascular and heart diseases in atomic bomb survivors: 1950–2003. Radiat. Environ. Biopyhs. 57, 2018. pp. 17–29. [https://doi.org/10.1007/s00411-017-0722-5]

Schütze H, Manning CD. Foundations of statistical natural language processing. Cambridge, Mass, MIT Press. 1999. p. 304. ISBN 978-0-262-13360-9.

Shuryak I, Fornace AJ, Datta K, Suman S, Kumar S, Sachs RK, Brenner DJ. Scaling human cancer risks from low LET to high LET when dose-effect relationships are complex. Rad. Res. 187, 2017. pp. 476–482.

Silverman BW. Density estimation for statistics and data analysis. London: Chapman & Hall/CRC. 1986: ISBN 978-0-412-24620-3. p. 45.

Simonsen LC, Slaba TC, Guida P, Rusek, A. NASA’s first ground-based galactic cosmic ray simulator: enabling a new era in space radiobiology research. PLOS Biology, 2020. [https://doi.org/10.1371/journal.pbio.3000669]

Simonsen LC, Slaba TC. Ensemble Methodologies for Astronaut Cancer Risk Assessment in the face of Large Uncertainties. NASA TP 2020-5008710, 2020.

Siranart N, Blakely EA, Cheng A, Handa N, Sachs RK. Mixed beam murine harderian gland tumorigenesis: predicted dose-effect relationships if neither synergism nor antagonism occurs. Rad. Res. 186, 2016. pp. 577–591.

Slaba TC, Qualls GD, Clowdsley MS, Blattnig SR, Walker SA, Simonsen LC. Utilization of CAM, CAF, MAX, and FAX for space radiation analyses using HZETRN. Adv. Space Res. 45, 2010. pp. 866–883.

Slaba TC, Blattnig SR, Norbury JW, Rusek A, La Tessa C. Reference field specification and preliminary beam selection strategy for accelerator-based GCR simulation. Life Sci. Space Res. 8, 2016. pp. 52–67.

Slaba TC, Bahadori, AA, Reddell, BD, Singleterry, RC, Clowdsley, MS, Blattnig, SR, Optimal shielding thickness for galactic cosmic ray environments. Life Sci. Space Res. 12; 2017. pp. 1–15.

Slaba TC, Whitman K. The Badhwar-O’Neill 2020 model. Space Weather 18, 2020. e2020SW002456.

Slaba TC, Wilson JW, Werneth CM, Whitman K. Updated deterministic radiation transport for future deep space missions. Life Sci. Space Res 27, 2020. pp 6–18.

Slingo J, Palmer T. Uncertainty in weather and climate prediction. Philos. Trans. A Math Phys. Eng. Sci. 369, 2011. pp. 4751–4767.

Smith T, Ross A, Maire N, Chitnis N, Studer A, Hardy D, et al. Ensemble modeling of the likely public health impact of a pre-erythrocytic malaria vaccine. PLoS Med 9, 2012. e1001157. [https://doi.org/10.1371/journal.pmed.1001157]

Tebaldi C, Knutti R. The use of the multi-model ensemble in probabilistic climate projections. Philos. Trans. A Math Phys. Eng. Sci. 365, 2007. pp. 2053–2075. [doi:10.1098/rsta.2007.2076]

Townsend LW, Nealy JE, Wilson JW, Simonsen LC. Estimates of galactic cosmic ray shielding requirements during solar minimum. NASA TM-4167, 1990.

Trani D, Datta K, Doiron K, Kallakury B, Fornace AJ. Enhanced intestinal tumor multiplicity and grade in vivo after HZE exposure: mouse models for space radiation risk estimates. Radiat. Environ. Biophys. 49, 2010. pp. 389–396.

UNSCEAR, United Nations Scientific Committee on the Effects of Atomic Radiation. Sources and effects of ionizing radiation. UNSCEAR 2006 report to the general assembly, with scientific annexes. United Nations, New York NY, 2006.

Walker SA, Townsend LW, Norbury JW. Heavy-ion contributions to organ dose equivalent for the 1977 galactic cosmic ray spectrum. Adv. Space Res. 51, 2013. pp. 1792–1799.

Walsh L, Schneider U, Fogtman A, Kausch C, McKenna-Lawlor S, Narici L, Ngo-Anh J, Reitz G, Sabatier L, Santin G, Sihver L, Straube U, Weber U, Durante M. Research plans in Europe for radiation health hazard assessment in exploratory space missions. Life Sci. Space Res. 21, 2019. pp. 73–82. [https://doi.org/10.1016/j.lssr.2019.04.002]

Werneth CM, Maung KM, Blattnig SR, Clowdsley MS, Townsend LW. Radiation shielding effectiveness with correlated uncertainties, Rad. Meas. 60, 2014. pp. 23–34.

Werneth CM, Xu X, Norman RB, Maung K, Relativistic three-dimensional lippmann-schwinger cross sections for space radiation applications. Nuc. Instr. Meth. Phys. Res. B 413; 2017. pp. 75–78.

Wilson JW, Cucinotta FA, Shinn JL. Cell kinetics and track structure. In: Swenberg CE, et al., Biological Effects and Physics of Solar and Galactic Cosmic Rays. Plenum Press. New York, NY, 1993. pp. 295–338.

Wilson JW, Slaba TC, Badavi FF, Reddell BD, Bahadori AA, Solar proton exposure of an ICRU sphere within a complex structure: combinatorial geometry. Life Sci. Space Res. 9; 2016. pp. 69–76.

Wilson JW, Werneth CM, Slaba TC, Badavi FF, Reddell BD, Bahadori AA, Effects of the Serber first step in 3DHZETRN-v2.1. Life Sci. Space Res. 26; 2020. pp. 10–27.

Yucker WR, Huston SL, The computerized anatomical female. Final Report, MDC-6107, McDonnell Douglas Company; 1990.

Yucker WR, Reck RJ, Computerized anatomical female body self-shielding distributions. Report, MDC 92H0749, McDonnell Douglas Company; 1992.

Zeitlin C, Hassler DM, Cucinotta FA, Ehresmann B, Wimmer-Schweingruber RF, Brinza DE. et al. Measurements of energetic particle radiation in transit to Mars on the Mars science laboratory. Science 340, 2013. pp. 1080–1084.

